# Genetic factors acting prior to dormancy in sour cherry influence bloom time the following spring

**DOI:** 10.1101/2023.11.09.566501

**Authors:** Charity Z. Goeckeritz, Chloe Grabb, Rebecca Grumet, Amy F. Iezzoni, Courtney A. Hollender

## Abstract

Bloom time is central to tree fruit production, and for *Prunus* species floral development leading up to bloom spans four seasons. Understanding this entire process is crucial for developing strategies to manipulate bloom time to prevent crop loss due to climate change. Here, we present a detailed examination of flower development from initiation until bloom for early- and late-blooming sour cherries (*Prunus cerasus*) from a population segregating for a major bloom time QTL on chromosome 4. Using a new staging system, we identified floral buds from early-blooming trees were persistently more advanced than those from late-blooming siblings. A gDNA coverage analysis revealed the late-blooming haplotype of this QTL, *k*, is located on a subgenome originating from the late-blooming *P. fruticosa* progenitor. Transcriptome analyses identified a large number of genes within this QTL as differentially expressed between early- and late-blooming trees during the vegetative-to-floral transition. From these, we identified candidate genes for the late bloom phenotype, including multiple transcription factors homologous to REproductive Meristem (REM) B3 domain-containing proteins. Additionally, we determined the basis of *k* in sour cherry is likely separate from candidate genes found in sweet cherry – suggesting several major regulators of bloom time are located on *Prunus* chromosome 4.

**HIGHLIGHT:** Dormancy is a main effector of bloom time in fruit trees. However, developmental, genetic, and transcriptomic analyses indicate differences in flower development before dormancy significantly influence flowering time in cherry.

## INTRODUCTION

*Prunus cerasus* L. (sour cherry) is an important temperate fruit tree in the Rosaceae family, which also contains sweet cherry, peach, apple, and pear (Xiang *et al*., 2017). Sour cherries are prized for their uniquely sweet and acidic flavor, and their processing qualities make them ideal for juice, jam, compote and pie. Apart from fruit quality, a critical trait for this industry is bloom time since it is essential for successful cross pollination and optimal fruit set. For instance, in 2012 a series of unexpectedly warm days in late March pushed fruit tree flower development far past their typical stage in Michigan at that time of year. Subsequently, a late spring freeze killed vulnerable flowers and annihilated over 90% of the expected sour cherry crop (Kistner *et al*., 2018). Unfortunately, this was not an isolated incident, as spring freezes caused significant losses in 2002, 2007, 2020 and 2021. Moreover, climate change is anticipated to increase the frequency of these events (Marino *et al*., 2011; Kistner *et al*., 2018; Unterberger *et al*., 2018). Therefore, strategies ranging from cultural practices to genetic improvement to control bloom time are sorely needed for a healthy and stable market.

Sour cherry is a recent allotetraploid with progenitors closely related to *Prunus avium* (sweet cherry) and *Prunus fruticosa* (ground cherry) (Olden and Nybom, 1968; Iezzoni and Hancock, 1984; Brettin *et al*., 2000; Bird *et al*., 2022). The recent publication of a reference genome for the cultivar Montmorency revealed three distinct subgenomes: A, A’, and B, with subgenome B at twice the dosage of A and A’ (Goeckeritz *et al*., 2023*b*). The two copies of subgenome B were inherited from the *P. avium*-like ancestor while subgenomes A and A’ originated from the *P. fruticosa*-like ancestor, which was also shown to be an allotetraploid of two unknown *Prunus* species. *Prunus avium* originates from the Mediterranean region south of the Black Sea, while *P. fruticosa* is typically found at more northern latitudes including Ukraine, Russia, and Kazakhstan (Quero-García *et al*., 2017). The latter shows some of the latest bloom times of the entire *Prunus* genus; in a common orchard, some *P. avium* accessions will bloom weeks earlier than *P. fruticosa*. Given their native habitats, this is consistent with observations in other species that show environmental signals such as photoperiod, temperature, and latitude have major implications for bloom time (Rohde *et al*., 2011; Hung *et al*., 2012; Hardigan *et al*., 2017). Comparatively, sour cherry genotypes exhibit diverse bloom times spanning the ranges of both progenitors. Therefore, fine-tuning bloom time using genetic approaches is a promising approach.

Bloom time is a highly heritable, quantitative trait that is a collection of many sequential developmental processes (Chardon *et al*., 2004; Salomé *et al*., 2011). A clear, comprehensive view of floral development from initiation to anthesis would be useful for designing new cultural practices to manipulate bloom time. Like other Rosaceous tree fruits, sour cherry begins its flower development in the summer (Diaz *et al*., 1981; Guimond *et al*., 1998). The apices in lateral and spur positions of the new year’s growth are vegetative at first but have the potential to become floral (Diaz *et al*., 1981). Although the molecular triggers for the floral switch are not fully understood, they have become clearer in the past decade. Several transcriptional studies and plant growth regulator applications have indicated that hormonal balance plays an important role in regulation of floral initiation in Rosaceous fruit trees. A low gibberellic acid (GA) to cytokinin (CK) ratio, as well increased abscisic acid (ABA) and indole-acetic acid (IAA; auxin) have been implicated to promote floral initiation in apple and sweet cherry (Li et al., 2018, 2019; Vimont *et al*., 2019; Villar *et al*., 2020; Bradley and Crane, 1960; Li *et al*., 2018; Zhang *et al*., 2019; Gottschalk *et al*., 2021). A few studies have shown variation in the timing of apple and peach floral development between years, suggesting photoperiod is not a strong environmental cue for initiation – at least for those genotypes (Raseira and Moore, 1987; McArtney *et al*., 2001; Kofler *et al*., 2019). However, physiological evidence indicates that there are strong temperature effects on this developmental process, with different species exhibiting distinct optimum temperatures for floral initiation (Sønsteby and Heide, 2019; Heide *et al*., 2020).

Floral organ development in rosaceous fruit trees initiates in the summer and involves the activity of MADS-box transcription factors including members of the *SHORT VEGETATIVE PHASE (SVP), SEPALLATA (SEP), PISTILLATA (PI)*, and *AGAMOUS (AG)* subfamilies (Villar *et al*., 2020). *APETALA* 2/*ETHYLENE RESPONSE FATOR* (*AP2/ERF*), *MYB*, *basic HELIX-LOOP-HELIX* (*bHLH*), and *NAC* transcriptions factors are also expressed in floral buds at this time (Vimont *et al*., 2019; Villar *et al*., 2020; Goeckeritz and Hollender, 2021). ABA and CK are at their highest concentrations around the time of floral induction but then decline as organogenesis proceeds; IAA concentrations show a similar trend in apple, but are reversed in sweet cherry (Li *et al*., 2018; Vimont *et al*., 2019; Villar *et al*., 2020). As day length and temperature declines, trees gradually enter a state of dormancy. For *Malus* and *Pyrus*, low temperatures appear to be the key environmental inducer of dormancy, while *Prunus* seems to monitor both temperature and photoperiod (Heide and Prestrud, 2005; Heide, 2008). Pioneering work in poplar exploring tree physiology during dormancy transitions has greatly furthered our understanding of this biological process. Decreasing day length causes the build-up of callose in phloem plasmodesmata, obstructing transport of growth-promoting signals to the apical meristem (Singh *et al*., 2018, 2019; Tylewicz *et al*., 2018; André *et al*., 2022).

Traditionally winter dormancy has been categorized into two discrete phases: endodormancy and ecodormancy (Lang *et al*., 1987). To relieve the growth inhibition wrought by endodormancy, chilling must be sensed by the quiescent meristem; for ecodormancy, warmth must return for growth resumption and budbreak. Both phases are under genetic control, although chilling, or length of endodormancy, has a much higher and more reliable broad sense heritability (Fan *et al*., 2010; Sánchez-Pérez *et al*., 2012; Allard *et al*., 2016; Kitamura *et al*., 2018; Branchereau *et al*., 2022). ‘Chill requirements’ describe the amount of cold a genotype must experience to reach 50% budbreak after some specified amount of time in forcing conditions, while ‘heat requirements’ refers to the growing degree units required once chilling needs are fulfilled. Abundant evidence demonstrates further chilling beyond the defined requirements increases heat’s effectiveness on development, and that historical estimations of chilling requirements are insufficient for full, synchronized bloom and commercial productivity (Okie and Blackburn, 2011; Luedeling, 2012; Campoy *et al*., 2019).

ABA is considered a dormancy maintenance hormone while GA appears to be involved in dormancy breakage through activation of genes essential to basic metabolism and growth (Zhuang *et al*., 2013, 2015; Yang *et al*., 2020). Transcriptomic data in a variety of Rosaceous fruit trees support this, and the antagonism between these two hormones in the context of dormancy is a key discussion point of many reviews (Cooke *et al*., 2012; Zhuang *et al*., 2013; Beauvieux *et al*., 2018; Falavigna *et al*., 2019; Vimont *et al*., 2019; Fadón *et al*., 2020; Yang *et al*., 2020; Goeckeritz and Hollender, 2021). Less studied but gaining traction is the proposed influence of reactive oxygen species (ROS) on the ABA:GA interplay, dormancy, and the cell cycle (Beauvieux *et al*., 2018). Indeed, at non-lethal levels, ROS such as H_2_O_2_ and O_2_ ^•-^ have been likened to hormones since they have potent effects on genes related to cell wall-loosening, expansion, and cycling (Sudawan *et al*., 2016; Ionescu *et al*., 2017; Velappan *et al*., 2017; Roussos, 2023). Several studies in perennials have shown ROS accumulation associates with deeper dormancy and steadily declines as meristems become more responsive to growth-promoting temperatures, a trend quite like ABA and opposite to GA (Wang *et al*., 2016; Lv *et al*., 2018; Yang *et al*., 2020; Sapkota *et al*., 2021; Wen *et al*., 2023).

Genetic analyses in peach resulted in the identification of six tandemly-arrayed homologs of SVP known as the Dormancy Associated MADs-box genes (*DAM*s; Bielenberg *et al*., 2004). These genes’ roles in *Prunus* dormancy transitions have been intensely studied through transcriptomic, epigenomic, and transgenic means (Leida *et al*., 2012; Bai *et al*., 2013; de la Fuente *et al*., 2015; Rothkegel *et al*., 2017; Prudencio *et al*., 2019; Zhu *et al*., 2020; Zhao *et al*., 2023). Although tandemly-arrayed, these genes appear to have distinct expression patterns, with several connected to higher chilling needs and later bloom times (Jimenez *et al*., 2010; Leida *et al*., 2012; Rothkegel *et al*., 2017; Calle *et al*., 2020; Zhao *et al*., 2023). Several findings have shown positive feedback loops between ABA levels and *DAM* gene expression (Singh *et al*., 2019; Yang *et al*., 2020; Zhao *et al*., 2023). In contrast, cytokinin signaling was linked with the downregulation of an apple *DAM1* ortholog and dormancy release (Cattani *et al*., 2020). Once the meristem accumulates sufficient chill and warmth returns the following spring, flowers grow exponentially until final bloom. GA, sugar transport, and genes involved in basic metabolic processes such as cell division and differentiation are all crucial to growth resumption in spring (Zhuang *et al*., 2015; Lv *et al*., 2018; Vimont *et al*., 2019).

In *Prunus*, the *DAM* genes have been independently attributed to chilling perception, dormancy transitions, and contrasting bloom times in Quantitative Trait Locus (QTL) mapping and other genetic association studies (Fan *et al*., 2010; Calle *et al*., 2020; Zhao *et al*., 2023). However, QTL have been found on all *Prunus* chromosomes, with large effect QTL that often overlap being found on chromosome 4 (Quilot *et al*., 2004; Dirlewanger *et al*., 2012; Ballester *et al*., 2001; Fan *et al*., 2010; Sánchez-Pérez *et al*., 2012; Cai *et al*., 2018; Kitamura *et al*., 2018; Branchereau *et al*., 2022). As *Prunus* genomes exhibit extensive synteny, future gene discoveries will likely expand our understanding of bloom time in sister species as well (Calle *et al*., 2020; Branchereau *et al*., 2022).

In the following work, we take a holistic approach to identify candidate genes in a population of sour cherry individuals segregating for bloom time for which a large (c. 8 Mb) QTL on linkage group 4 named *qP-BD4.1^m^* explains 27.9% of phenotypic variation (Cai *et al*., 2018). Previous marker phasing led to the identification of seven haplotypes of this QTL. One haplotype in particular, *k*, delays average bloom time in an additive manner by approximately three days. Each tetraploid parent contained one copy of *k* – thus, F1 progeny inheriting 2*k* were more likely to be late-blooming (LB) while those inheriting 0*k* were more likely to be early-blooming (EB). We assessed floral biology in 2*k* and 0*k* full-siblings from the time of floral initiation through anthesis to understand when and how these individuals differed in their developmental trajectories. The comprehensive picture of flowering phenology in these individuals led to the creation of a floral development staging system that will be easily adaptable for other *Prunus* species. We also identified genetic variants unique to the *k* haplotype and conducted differential gene expression and co-expression network analyses to select high-quality candidate genes underlying the late-blooming trait.

## MATERIALS AND METHODS

### Sampling procedures for morphological, physiological, gDNA-sequencing, and RNA-sequencing analyses

All materials were collected from trees at Michigan State University’s (MSU) Clarksville Research Center (42.87361004512462, -85.25873447306927). The number of individuals included in observational experiments from year to year depended on tree health, size, and available time and resources. Tier, row, and tree numbers are given in **Table S1**, alongside their identity in the present work. Whole apices/flower buds for tissue fixation and sectioning, pistil growth measurements, and ROS extraction were sampled randomly on the east and west sides of the trees. Branches for forcing experiments were also collected randomly but with the additional goal of having at least 25 flower buds total per genotype per collection date from at least two separate branches. We had high confidence that apices/buds collected were or would become flowers since *Prunus* species tend to flower and fruit in consistent positions from year to year (Diaz *et al*., 1981). Replicates intended for ROS quantification were a pool of four adjacent flower buds. For RNA-sequencing, three biological replicates for six trees (3 EB and 3 LB) were sampled randomly as a pool of 6 neighboring apices at three dates (12 June 2019, 29 June 2019, 18 July 2019). Sampling occurred at approximately the same time of day for each collection (12:00 – 14:00) to mitigate gene expression noise between time points. Whole apices were excised with a razor blade, immediately flash-frozen in liquid nitrogen in the field, then stored at -80°C until RNA extraction. Dates for pistil and histological analyses, branches for chill assessments, flower buds for ROS (H_2_O_2_) quantification, and tissue collection for RNA-sequencing are given in **Tables S2 – S5**, with corresponding growing degree days (GDDs) and Dynamic Model chill portions (where applicable). For the present study, growing degree hours / days (GDHs / GDDs) were measured using the method developed by Richardson et al. (1975):

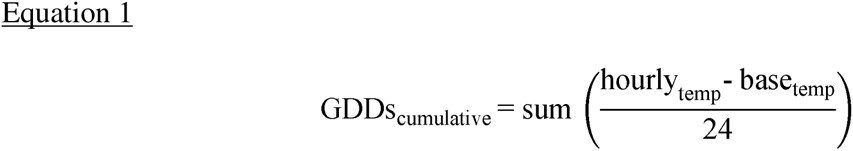

with a base temperature of 4.4 °C and no maximum temperature limit (Richardson *et al*., 1974). Chill portions were calculated using chillR (Luedeling *et al*., 2013). All hourly temperature data was downloaded from the Clarksville, MI, MSU weather station: https://legacy.enviroweather.msu.edu/weather.php?stn=clr (2023). *Tissue fixation, embedding, histological sectioning, and staging*

Fixation, embedding and sectioning were done using the protocol described by Goeckeritz et al. (2023*a*). Briefly, flowers were collected in the field and either placed in a 2.5% glutaraldehyde / 2.5% formaldehyde / 0.1 M sodium cacodylate fixative solution (pH 7.2) after excising the tip of the bud to maximize infiltration, or placed in fixative after bud scales were removed in the lab. They were then subject to at least two 30 min vacuum infiltrations before being stored in fixative at 4 °C until dehydration and embedding. Sectioning was done at a 1 µm thickness using a PMC XL Ultramicrotome (Boeckeler Instruments, Tucson, AZ, USA), and sections were heat-mounted to glass slides and stained with a diluted toluidine blue solution (catalog no. 14950, Electron Microscopy Sciences).

The staging system for *Prunus cerasus* was created by studying histological patterns in development among individuals throughout two seasons (2018-19; 2019-20) and a review of the literature (Endress, 2015; Becker, 2020). Features such as the emergence of organ primordia (sepals, petals, stamens, pistil) and the emergence of specific cell layers were used to demarcate staging. Particular attention was paid to the male (anthers) and female (ovule) gametophytes since their development was more easily defined by discrete stages; in contrast, sepal and petal advancement mainly manifested as relatively subtle cell divisions and elongation. SEM photos of related species’ development (*P. laurocerasus* and *P. serotina*) assisted with 3-dimensional visualization of the pistil/carpel and was critical to determining the approximate timing of closure (Wang *et al*., 2019). Illustrations in Esau 1965, Bierhorst 1971, Foster and Gifford 1974, Mauseth 1988, and Fahn 1990 were fundamental in accurately recognizing features of micro-and megagametogenesis; for example, anther cell layers and the general progression of the embryo sac. Stages were hand drawn using an iPad Air (5^th^ generation) and the ProCreate version 5.3.5 software.

Unpaired, two-sample Wilcoxon rank sum tests (equivalent to two-sample Mann-Whitney U tests) were performed to identify statistically significant differences in development between EBs and LBs for nine pre-dormancy dates. The response variable was treated as numeric and each flower bud was considered an independent sample for its respective bloom group, irrespective of the individual tree it originated from. P-values for the nine tests were corrected using a Bonferroni calculation.

### Pistil measurements and analysis

Pistil measurements for trees were done for years 2018-2019 (three trees per bloom group), 2019-2020 (six EB, four LB), and 2021-2022 (seven EB, five LB) as growing degree days accumulated. 5 – 10 flowers from separate buds per tree were dissected using a Nikon SMZ800N stereo microscope and imaged with a Nikon DS-Fi3 color camera and Nikon NIS-Elements BR 4.60.00 software (Nikon, Tokyo, Japan). Pistils were measured from the tip of the stigma to the ovary base (see **Figure 10** inset). On the day of bloom (approximately 50% open flowers on the tree), 30 open flowers were collected for each genotype to normalize measurements according to the final pistil length of each individual; thus, the data is expressed as a fraction of final pistil length. Due to the different numbers of individuals and collection times per year, each year was modeled separately as:

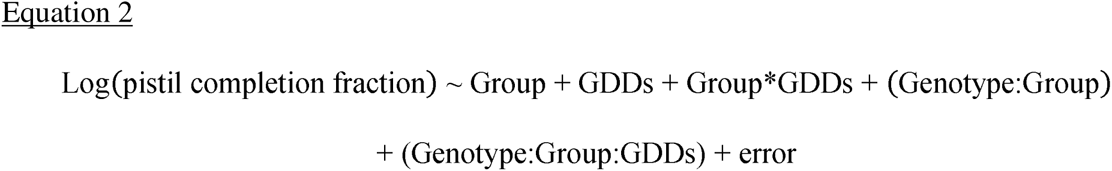

A logarithmic transformation was necessary to approach statistical normality. ‘Group’ was a fixed effect factor with early- and late-blooming levels and ‘GDDs’ was a fixed, ordered factor (to detect significant differences at specific dates) with as many levels as collection time points that year. ‘Group * GDDs’ represented the developmental effect of the interaction of bloom group with accumulating heat. The ‘(1 | Genotype:Group)’ represented the random effects that each individual tree, which was nested within bloom group, contributed to development. Lastly, the ‘(1 | Genotype:Group:GDDs)’ term represented the random effect on development that each tree had at each collection time point. The raw data (pistil measurements / pistil completion fraction) is provided as **Files S1 – S3** and R scripts used to analyze the data and create the figures are available at https://github.com/goeckeritz/sourcherry_bloomtime.

### Forcing experiments and analysis

Forcing experiments took place in the dormant seasons of 2019-2020 and 2021-2022. Branches for forcing were collected from six EB and three LB individuals in 2019-2020 at eight dates (26.6 – 91.5 CP), and from 6 – 7 EB and 3 – 4 LB individuals in 2021-2022 at nine dates (20.7 – 67.7 CP). Budbreak scoring was done using a slightly modified BBCH scale, a decimal coding system describing the phenology of woody perennials (Fadón *et al*., 2015). The dependent variable for each statistical model was the proportion of flowers reaching BBCH stages 51, 52, 53, and 54 (i.e., four statistical models were created per year); data after stage 54 is not presented since the health of the branches seemingly declined after this developmental stage. The ends of each collected branch were given fresh cuts, placed in distilled (DI) water, and forced out of dormancy in a chamber held at a constant 22°C (+/- less than 1 °C) with fluorescent lighting set to a 12-hour photoperiod. Branches received fresh cuts and DI water thereafter every 1 – 2 days, when scoring of each flower bud occurred. Briefly, stage 50 was described as dormant with no visible bud swelling. Stage 51 was characterized by first swelling, with some evident green tissue emerging from the breaking bud. Stage 52 was added to the scoring system as an intermediate between stages 51 and 53. A flower bud was considered stage 53 when green tissue could be seen at least halfway down opposite sides of the bud. Stage 54 was reached when the bud cracked open due to the enlargement of the flowers inside. If any field GDDs had accumulated after January 1^st^ in the year of study, these GDDs were added at the start of the forcing experiment for that collection date (e.g., for the collection on 10 Jan 2020, 88.1 GDHs, or 3.7 GDDs, had accumulated since 1 Jan 2020). However, this value was negligible as field heat accumulation was no higher than 178.2 GDHs / 7.4 GDDs for any collection after 1 Jan for both years. Chill was calculated using the Dynamic Model in the chillR package starting 1 October (Fishman *et al*., 1987*a*,*b*; Luedeling *et al*., 2013; R Core Team, 2022).

The two seasons were analyzed independently since there was not complete overlap in the accessions and chill portions at each collection date between years. For 2019-2020, 5 early-blooming genotypes (with 0*k*) and 3 late-blooming genotypes (with 2*k*) were scored and analyzed. For 2021-2020, 7 early-blooming genotypes (5 were common to both years) and 4 late-blooming genotypes (3 were common to both years) were scored and analyzed. For each year, separate statistical models, including both fixed and random effects, were constructed for BBCH stages 51, 52, 53, and 54. Models were not created for later stages due to concerns about branch health and thus, reliability of the data. Proportion of flowers at or above a particular stage (*X*) were modeled as such:

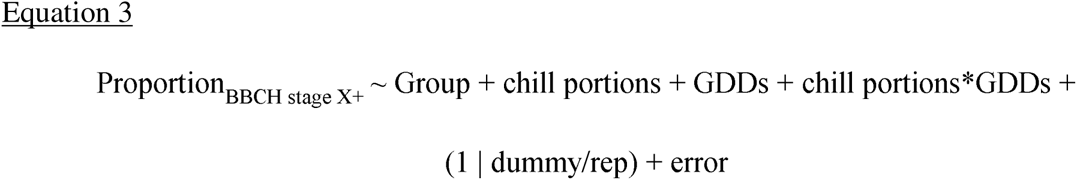

where ‘Group’ was either early- or late-blooming (0*k* or 2*k*, respectively), and ‘chill portions’ was the amount of chill accumulated between Oct 1^st^ and the collection date according to the Dynamic Model. ‘Chill portions’ was coded as an ordered factor as it was equivalent with collection date. GDDs was a continuous variable. To account for the repeated measurements within each collection, a dummy variable was first coded as a combination of an individual tree within each timepoint. Then, using the ‘corSymm’ function of R package ‘nlme’, the correlation structure was modeled by nesting the repeated measurement within this dummy variable (Pinheiro and Bates, 2023). Raw data for the two seasons of forcing experiments are given as **Files S4 – S5**, and R scripts for data analysis and figure generation can be found at https://github.com/goeckeritz/sourcherry_bloomtime.

### Measuring H_2_O_2_ content in early- and late-blooming individuals during dormancy transitions

We tried several fluorometric / spectrometric methods to measure hydrogen peroxide: including one based on Amplex Red™ (Chakraborty *et al*., 2016), 3-methyl-2- benzothiazolinone hydrazone and 3-(dimethylamino) benzoic acid (MBTH-DMAB) (Ngo and Lenhoff, 1980; Noctor *et al*., 2016), and xylenol orange (Cheeseman *et al*., 2006). Each method was subject to two control tests. In the first test, samples were measured at 5 - 30 min intervals for at least a half hour to ensure stable signal and therefore, stable concentration of H_2_O_2_. In the second, known concentrations of H_2_O_2_ were added to the sample extract and the correlation between the expected and observed quantities of H_2_O_2_ were noted. Only an adaptation of the xylenol orange method (for measurement in a 96-well plate) passed both controls, but it was sensitive to temperature and humidity. This may have been due to the solubility of KCN, a component of the extraction buffer. Therefore, the plate (Greiner Bio-One, Ref 655903) was pre-cooled prior to sample loading and kept on ice at all other times, being careful not to smudge the underside as this would affect signal readings. Additionally, all reagents and sample extracts were constantly kept on ice and all centrifugation steps were performed at 4° C. The humidity was monitored with a dehumidifier and kept under 30% during extraction to prevent condensation buildup on liquid N_2_ chilled mortars and pestles. Periodically during the dormant season (October – March; **Table S5**), at least three biological replications of four adjacent flower buds were collected per tree and immediately frozen in liquid nitrogen. Flower buds of four late-blooming and six early-blooming individuals were collected at all five time points. Given reports that ROS degrade over time even when materials are stored at -80° C, all samples were kept in liquid nitrogen in a large Dewar until extraction (Cheeseman, 2006).

On the day of extraction, the fresh weight of each sample (four flower buds) was determined by first measuring each tube containing 5 mL extraction buffer without sample, weighing the tube again after adding ground sample material (thawed), and calculating the difference. Samples were ground to a fine powder using a mortar and pestle and quickly added to 5 mL of cold NaH_2_PO_4_ buffer at pH 6.4 + 5 mM KCN, which was then vortexed immediately. Eight fresh H_2_O_2_ standards between concentrations of 0 – 4 µM were prepared alongside samples by serial diluting a 30% H_2_O_2_ solution (Thermo Scientific, H325-500) in extraction buffer. Vortexed samples were then centrifuged at 4° C at max speed for 10 minutes. Meanwhile, a working solution was prepared containing 500 µM ferrous (II) ammonium sulphate, 200 µM sorbitol, 50 mM H_2_SO_4_, 2% ethanol and 200 µM of xylenol orange. A ‘pre-working solution,’ without xylenol orange, was stored at room temperature; xylenol orange was not added until the day of extraction. After centrifuging samples, the supernatants were transferred to new tubes on ice. The chilled 96-well plate was then loaded by first adding 100 µL of the working solution to the empty wells. Then, 100 µL of the standards and samples were added to the wells (for a total volume of 200 µL). Each standard and sample had four technical replicates. After loading, the plate was placed back in a 4° C fridge and allowed to incubate for 60 – 90 minutes (the reaction is slower in the cold). Multiple readings of the plate were sometimes warranted if the standard curve had not yet linearized, indicating the reaction was incomplete. The plate was then removed from the fridge and allowed to sit at room temperature for 30 seconds, the bottom of the plate was checked to contain no condensation, and then absorption was measured at 550 and 800 nM using a Synergy H1 microplate reader (BioTek Instruments, Inc). The data was exported to an Excel spreadsheet with Gen5 software version 2.01.14 (BioTek Instruments, Inc) and µmoles of H_2_O_2_ per gram of fresh weight for every well were determined using the standard curve. In 4 of 14 cases, the 0 µM standard was dropped since it significantly skewed the curve. Standard curve r^2^ values ranged from 0.86 – 0.99. After all extractions, there were 2-3 biological replications for each tree per time point. The detailed extraction protocol can be found at https://github.com/goeckeritz/sourcherry_bloomtime.

The raw data was first filtered by dropping negative values (representing readings where background noise was higher than the signal from the sample). Then the data was modeled as:

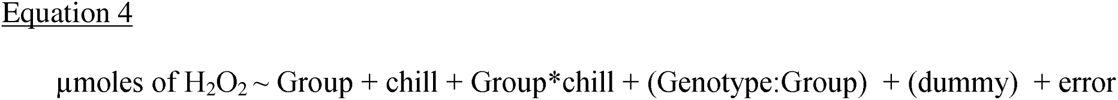

Where the fixed factor ‘Group’ contained levels early- and late-blooming, and the fixed factor ‘chill’ described how much field chill had accumulated since October 1^st^, 2019 (synonymous with collection date). ‘(1 | Genotype:Group)’ represented the random variation introduced by individual genotypes within a bloom group, and ‘(1 | dummy)’ represented the technical replicate variation, which was a combination of repeated measurements nested within a plate, genotype, and collection date (chill level). The data for these analyses are included in **File S6**.

### Manual curation of gene models within the chr4 QTL region

Prior to predicting genetic variation effects on gene models within the chr4 QTL on subgenome A (pipeline subsequently explained), the genes within this region were manually examined against the RNAseq data from the current study using Apollo v 2.6.5 (Dunn *et al*., 2019). The RNAseq reads were aligned to ‘Montmorency’ subgenome A as described below. Manual curation for these genes was done to refine their quality for more accurate candidate gene selection.

### Genome sequencing, coverage assessments, SNP-calling, and structural variant analysis

The ‘Montmorency’ reference genome contains phased subgenomes A, A’ and B, and recent findings suggest diverse sour cherries contain variable dosages of these subgenomes (Iezzoni and Rhoades, personal communication). All seven haplotypes identified at the chr4 QTL (*k, a, h, n, o, i,* and *g*) from the previous work (Cai *et al*., 2018) were included amongst the sequenced individuals. We examined the relative read coverage of each subgenome within the QTL for the 12 individuals alongside their parents, ‘Balaton’ and ‘Surefire’ (**Figure 13**). Logical statements comparing haplotypes and subgenome composition amongst individuals were developed to identify which haplotype originated from which subgenome (described fully in **File S7**).

DNA of young, unexpanded leaves was extracted from 14 individuals (7 early-blooming, 5 late-blooming, and the two parents) with a DNeasy Mini Plant Kit (Qiagen, Hilden, Germany) and quantified with a Quant-iT^TM^ PicoGreen^TM^ dsDNA assay kit (Thermo Fisher Scientific, Waltham, MA) or using the Qubit DNA Broad Range kit and Qubit 4 Fluorometer (Thermo Fisher Scientific Inc, Waltham, MA). Quality was assessed with an Agilent 4200 TapeStation HS DNA1000 and Kapa Biosystems Illumina Library Quantification qPCR assays. Illumina TruSeq Nano DNA 150 bp paired-end (PE) libraries were prepared at MSU’s Research Technology Support Facility (RTSF) and sequenced at 20X (progeny) or 60X (parents) using a HiSeq4000 instrument. At least 22.6 million paired end reads were produced for each sample and checked for quality using FASTQC (Andrews, 2010). Reads were then trimmed using Trimmomatic v 0.39 and aligned to the scaffolded ‘Montmorency’ reference genome (including subgenomes A, A’, and B) using BWA v 0.7.17 (Li and Durbin, 2009). The resulting .bam files were sorted, and duplicates were removed with picard tools v 2.25.0 (‘PicardTools’, 2018). To produce the coverage plots, read depth was calculated after alignment to the reference genome for every position using SAMtools v 1.11 (Li *et al*., 2009). Then, a custom bash loop was written to average the read depth over every 500 kb window for all 14 samples (see coverage_loops.sh, https://github.com/goeckeritz/sourcherry_bloomtime). These files were plotted in R using ggplot2 to visualize and determine the subgenome dosage at the bloom time locus (**Figure 13**; (R Core Team, 2022; Wickham).

Since *k* was determined to be located on subgenome A, we excluded Early 1 (subgenome composition A’BBB) from SNP and structural variant calling. SNP calls were made only in the QTL region (Pcer_chr4A: 8324991- 16386209), and ploidy was adjusted for every individual according to their subgenome A dosage. This required two separate regions to be called for ‘Surefire’ and Early 7, as their dosage increased (due to the *g* haplotype) at approximately 12583308 bp along the length of chromosome 4 (determined by examining depth files). We used HaplotypeCaller within GATK v 4.2.0.0 and a pedigree file to call SNPs for all samples (Mckenna *et al*., 2010). Next, CombineGVCFs was used to merge the two parts of ‘Surefire’ and Early 7; then, these complete files were combined with all others individuals’ gvcf files. The variants were then hard-filtered based on quality metrics (see variant_filtration.sh, https://github.com/goeckeritz/sourcherry_bloomtime) and assigned a predicted effect on the gene model with snpEff (Cingolani *et al*., 2012). VariantsToTable was used to convert this file to a table, which was read into R for further filtering based on known inheritance patterns. For example, variants of interest include those that are unique to the individuals with 2*k* (late-blooming); and, if *k* is an individual’s only source of subgenome A (i.e., the individual is 2X subgenome A and does not have haplotype *i* or *g* in addition to 2*k*), the genotype calls between late-blooming siblings should be identical. In this situation, each late-blooming individual will have two alleles (e.g., T/T, A/A, C/T, etc.), one of which should come from ‘Balaton’ and the other from ‘Surefire.’ Using such logic, we further filtered the dataset (see further_filtering_and_plotting.R). Regular expressions were then used to extract lists of genes with different classifications of variants (e.g., high effect mutations, 5’UTR, upstream, etc.) from the filtered file (provided in **File S8**).

To identify putative structural variants upstream and within protein-coding genes, reads mapped to the chr4A QTL were pooled so that higher coverage would represent both the *k* and *i* haplotypes. These pools were genotyped alongside ‘Surefire’, which was sequenced to approximately 60X (15X per allele), and Early 7 (20X, 5X per allele). Surefire represented haplotypes *i* and *k* in the first half of the QTL and haplotypes *i*, *k*, and *g* in the latter half. Early 7 was genotyped separately for another representative of the *g* haplotype. Variants were called with Delly v 0.7.8 (Rausch *et al*., 2012). Limited coverage representing the *g* haplotype meant fewer variants were called compared to *i* and *k*, and the reads representing *g* in ‘Surefire’ were confounded with *i* and *k*. Nonetheless, several criteria were imposed to filter variants. First, *k*’s unique genotype calls were anticipated to be homozygous and different from *i* calls, which were also expected to be homozygous (only bi-allelic sites were examined). ‘Surefire’ was useful when *i* calls were missing in the first half of the QTL, because homozygous calls in ‘Surefire’ meant *i* and *k* were identical at the location in question and such a variant could be ruled out as causative for late bloom. The resulting 169 structural variants is a liberal estimate since in many instances, calls for haplotypes other than *k* were missing. To compensate for this, called variants predicted to affect DEGs of interest in **Table 1** were manually verified in IGV v 2.13.0 by examining coverage of individuals with *i, k*, and *g* haplotypes (Thorvaldsdóttir *et al*., 2013).

**Table 1:**
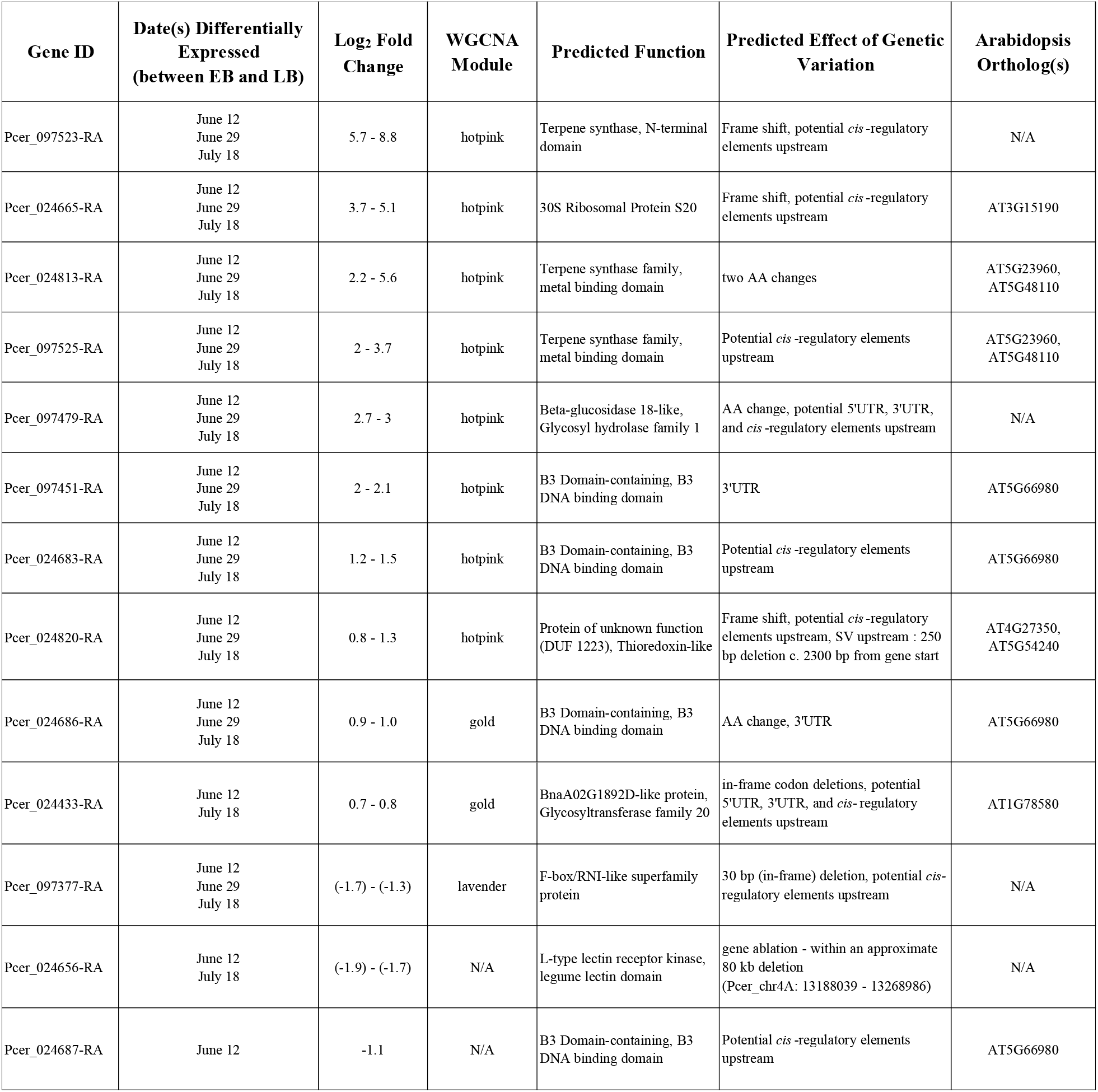
High-quality candidate genes for the basis of the *k* late-bloom phenotype. These contain genetic variation and are in most cases expressed in modules of interest. Genes beginning with 097 were manually edited with Apollo and thus are not indicative of relative chromosome position. Arabidopsis orthologs were identified using OrthoFinder v2.5.4.

| Gene ID | Date(s) Differentially Expressed<br>(between EB and LB) | Log<sub>2</sub> Fold Change | WGCNA Module | Predicted Function | Predicted Effect of Genetic Variation | Arabidopsis Ortholog(s) |
| --- | --- | --- | --- | --- | --- | --- |
| Pcer_097523-RA | June 12<br>June 29<br>July 18 | 5.7 - 8.8 | hotpink | Terpene synthase, N-terminal domain | Frame shift, potential *cis*-regulatory elements upstream | N/A |
| Pcer_024665-RA | June 12<br>June 29<br>July 18 | 3.7 - 5.1 | hotpink | 30S Ribosomal Protein S20 | Frame shift, potential *cis*-regulatory elements upstream | AT3G15190 |
| Pcer_024813-RA | June 12<br>June 29<br>July 18 | 2.2 - 5.6 | hotpink | Terpene synthase family, metal binding domain | two AA changes | AT5G23960,<br>AT5G48110 |
| Pcer_097525-RA | June 12<br>June 29<br>July 18 | 2 - 3.7 | hotpink | Terpene synthase family, metal binding domain | Potential *cis*-regulatory elements upstream | AT5G23960,<br>AT5G48110 |
| Pcer_097479-RA | June 12<br>June 29<br>July 18 | 2.7 - 3 | hotpink | Beta-glucosidase 18-like, Glycosyl hydrolase family 1 | AA change, potential 5'UTR, 3'UTR, and *cis*-regulatory elements upstream | N/A |
| Pcer_097451-RA | June 12<br>June 29<br>July 18 | 2 - 2.1 | hotpink | B3 Domain-containing, B3 DNA binding domain | 3'UTR | AT5G66980 |
| Pcer_024683-RA | June 12<br>June 29<br>July 18 | 1.2 - 1.5 | hotpink | B3 Domain-containing, B3 DNA binding domain | Potential *cis*-regulatory elements upstream | AT5G66980 |
| Pcer_024820-RA | June 12<br>June 29<br>July 18 | 0.8 - 1.3 | hotpink | Protein of unknown function (DUF 1223), Thioredoxin-like | Frame shift, potential *cis*-regulatory elements upstream, SV upstream : 250 bp deletion c. 2300 bp from gene start | AT4G27350,<br>AT5G54240 |
| Pcer_024686-RA | June 12<br>June 29<br>July 18 | 0.9 - 1.0 | gold | B3 Domain-containing, B3 DNA binding domain | AA change, 3'UTR | AT5G66980 |
| Pcer_024433-RA | June 12<br>July 18 | 0.7 - 0.8 | gold | BnaA02G1892D-like protein, Glycosyltransferase family 20 | in-frame codon deletions, potential 5'UTR, 3'UTR, and *cis*-regulatory elements upstream | AT1G78580 |
| Pcer_097377-RA | June 12<br>June 29<br>July 18 | (-1.7) - (-1.3) | lavender | F-box/RNI-like superfamily protein | 30 bp (in-frame) deletion, potential *cis*-regulatory elements upstream | N/A |
| Pcer_024656-RA | June 12<br>July 18 | (-1.9) - (-1.7) | N/A | L-type lectin receptor kinase, legume lectin domain | gene ablation - within an approximate 80 kb deletion (Pcer_chr4A: 13188039 - 13268986) | N/A |
| Pcer_024687-RA | June 12 | -1.1 | N/A | B3 Domain-containing, B3 DNA binding domain | Potential *cis*-regulatory elements upstream | AT5G66980 |

### RNA sequencing, differential gene expression, and weighted gene correlation network analyses

RNA from summer apices was extracted using a previously-developed protocol treated with a DNA-*free*^TM^ Kit (Thermo Fisher Scientific, Waltham, MA) to remove DNA, and assessed for initial quality using a Nanodrop One^C^ (Thermo Fisher Scientific, Waltham, MA) (Gasic *et al*., 2004). If purity was insufficient (260/230 < 1.8), samples were further cleaned and concentrated using the RNA Clean & Concentrator^TM^-25 kit (Zymo Research, Seattle, WA). Concentration of each sample was measured using the Qubit RNA Broad Range kit and Qubit 4 Fluorometer (Thermo Fisher Scientific Inc, Waltham, MA), and quality was assessed with an Agilent High Sensitivity RNA ScreenTape System (Agilent Technologies, Santa Clara, CA).

Illumina stranded mRNA 150 bp PE libraries were prepared at MSU’s RTSF before they were pooled and sequenced on approximately 0.8 lanes of a NovaSeq 6000, producing at least 39.8 million reads per library. A total of 48 libraries were sequenced: three replicates per tree for 12 June, two replicates per tree for 29 June, and three replicates per tree for 18 July. The same three trees were used as representatives of each bloom group (EB and LB) for each collection date (i.e., six trees were used).

Each resulting fastq file was confirmed to be of high quality with FASTQC. Reads were processed with Trimmomatic v 0.39, then aligned with STAR v 2.7.3a to *P. cerasus* ‘Montmorency’ subgenome A since the coverage assessment results suggested the gene(s) responsible for *k*’s late bloom phenotype were unique alleles of subgenome A (Dobin *et al*., 2013; Bolger *et al*., 2014). We were reassured that reads from subgenome B and A’ would align to orthologous genes on subgenome A since *Prunus* genomes are highly syntenic and collinear, and because alignment parameters were relaxed to allow for expected mismatches while still retaining sufficient mapping quality. Briefly, parameters --outFilterScoreMinOverLread (allowable ratio of mismatches to mapped length) --outFilterMatchNminOverLread, (minimum requirement of matched bases, normalized by read length) and --outFilterMatchNmin (minimum requirement of matched bases) were all set to zero to ensure maximum mapping. No more than 30% of each read’s mapped length was allowed to mismatch (--outFilterMismatchNoverLmax). These parameters had the potential of keeping very short (and potentially inaccurate) mappings; therefore, we checked the average input read length versus average mapped read length. The average input read length was 294 and above for each library (2 x 150 paired-end reads), and the lowest average mapped length was 251; thus, on average, at least 85% of each read’s full length was successfully mapped. Uniquely-mapped reads ranged from 87 – 92% per library, and there was no obvious mapping bias among certain trees or time points.

These alignments were used with ‘Montmorency’ subgenome A gene models (.gff3) to assess the expression of each gene. Expression was determined using StringTie v 2.1.3 and given as counts for the required input for the DESeq2 R package, and Fragments Per Kilobase per transcript per Million mapped reads (FPKM) as the required input for the WGCNA (Weighted Gene Correlation Network Analysis) R package (Langfelder and Horvath, 2008; Love *et al*., 2014; Kovaka *et al*., 2019; R Core Team, 2022).

For the differential expression (DE) analysis using DESeq2, transcripts with no expression across all samples were dropped from the analysis. Using a regularized log transformation, normalized expression counts for each library were visualized by a Principal Component Analysis (PCA). Two of the 48 samples were identified as outliers (27_04_34_6_12_19_R2 and 27_03_46_6_29_19_R3) and removed from further analysis. Counts were normalized and modeled using the DESeqDataSetFromMatrix function with the following independent variables: bloom group (early or late / 0*k* or 2*k*), the individual nested within the bloom group, date, and the interaction of bloom and date. Seven comparisons were performed to assess gene expression changes: **1)** early_v_late_Jun12, **2)** early_v_late_Jun29, **3)** early_v_late_Jul18, **4)** early_Jun12_Jun29, **5)** early_Jun29_Jul18, **6)** late_Jun12_Jun29, and **7)** late_Jun29_Jul18. These comparisons correspond to gene expression changes between early- and late-bloomers on 12 June, 29 June, and 18 July, then early-bloomers from 12 June to 29 June and 29 June to 18 July, and late-bloomers from 12 June to 29 June and 29 June to 18 July, respectively. As bloom time is a quantitative trait, even subtle changes in gene expression could be responsible for the phenotypic differences; therefore, genes with an absolute log_2_ fold change of 0.5 or higher and a p-value less than 0.05 were considered differentially expressed genes (DEGs). Genome-wide results and the subset of results within the QTL region on chromosome 4 can be found in **File S9**. Venn diagrams were created using R package VennDiagram (Boutros, 2022).

For network analyses, genes with an FPKM value < 5 across all samples were excluded. Log-transformed FPKM values were further filtered to remove genes that varied little between samples (coefficient of variation < 0.2). After hierarchical clustering, the two samples identified as outliers during the differential expression analysis using DESeq2 were again removed from the dataset prior to weighted gene correlation network analysis. A signed adjacency matrix with a power threshold of 14 was created and used to calculate the topological overlap dissimilarity matrix. This matrix was used to create a gene tree, which was used as input for cutreeDynamic. DeepSplit was set to 2, the minClusterSize (minimum number of genes in each module) was set to 30, and the cut height was 0.996. Next, the eigengenes were clustered to identify modules with similar expression patterns and modules below cutHeight < 0.20 were merged. The eigengene expression for each of the final modules was then plotted by date to identify patterns associating with bloom group (**Figure 17**).

### GO database creation and enrichment tests

GO (gene ontology) IDs for each ‘Montmorency’ subgenome A gene were determined with InterProScan v 5.33-72.0 as previously described and BLAST+ v 2.9 (Quevillon *et al*., 2005; Camacho *et al*., 2009). We formatted a BLAST+ database for the Arabidopsis proteins, and these were used as a query while sour cherry protein sequences were used as the target (Carbon *et al*., 2021). Only hits with p-value < 1.0 x 10^-10^ were kept and all other parameters were set to default. Arabidopsis GO information was downloaded from arabidopsis.org and a simple R script was used to merge the InterProScan, arabidopsis to sour cherry blast hits, and arabidopsis GO terms to create the final database (available upon request as the file was too large to upload to https://github.com/goeckeritz/sourcherry_bloomtime).

topGO was used to determine which biological processes (BP) were enriched in the differentially expressed genes and WGCNA modules (Alexa and Rahnenfuhrer, 2023). topGOfunctions.R is a function wrapper created previously by Mansfeld *et al*. 2020. Since the dds object from the differential expression analysis with DESeq2 was used as input for all GO enrichment tests, R code performing both the DE and WGCNA module tests can be found at the end of the DESeq2 R script. Using the ‘weight01’ algorithm, terms were considered significant if the Fisher test had a p-value < 0.05 (Fisher, 1954). Minimum node size was set to 50. The results of these tests can be found in **File S10** (DE analysis) and **File S11** (WGCNA modules).

### Comparing Regina QTL syntelogs

A syntenic analysis between ‘Regina’ chromosome 4 and ‘Montmorency’ chromosome 4A was conducted using MCScan, and the sour cherry syntelogs of genes within the sweet cherry QTL region were used as anchors to identify this region in sour cherry (Branchereau *et al*., 2022). All genes within this region were searched for differential expression between early- and late-blooming individuals for any sampling date in the current study.

### Identifying Arabidopsis Orthologs of Candidate Genes

OrthoFinder v 2.5.4 was run with default parameters to identify orthogroups of arabidopsis and *P. cerasus* subgenome A, A’, and B proteins (Emms and Kelly, 2019). Arabidopsis peptides were downloaded in October 2023 from arabidopsis.org (Araport11_pep_20220914.gz), and subgenome A, A’, and B peptide sequences were retrieved from the modified ‘Montmorency’ annotation where the gene models within the subgenome A QTL had been manually curated (available at https://github.com/goeckeritz/sourcherry_bloomtime). Arabidopsis ortholog(s) are listed in **Table 1** for each respective *P. cerasus* gene.

## RESULTS

The average range in bloom times between individuals of the segregating population with 0*k* and 2*k* was 72 GDDs and 10 days for the three years of this study. Three representative early-bloomers (0*k*) and three late-bloomers (2*k*) photographed on the same day in two of these years are shown in **Figure 1**. Unique identifiers for each tree along with bloom times (50% open flowers) in calendar days and Growing Degree Days (GDDs) in the years they were scored are given in **Table S1**. Materials collected and corresponding dates are fully documented in **Tables S2 – S5**.

**Figure 1:**
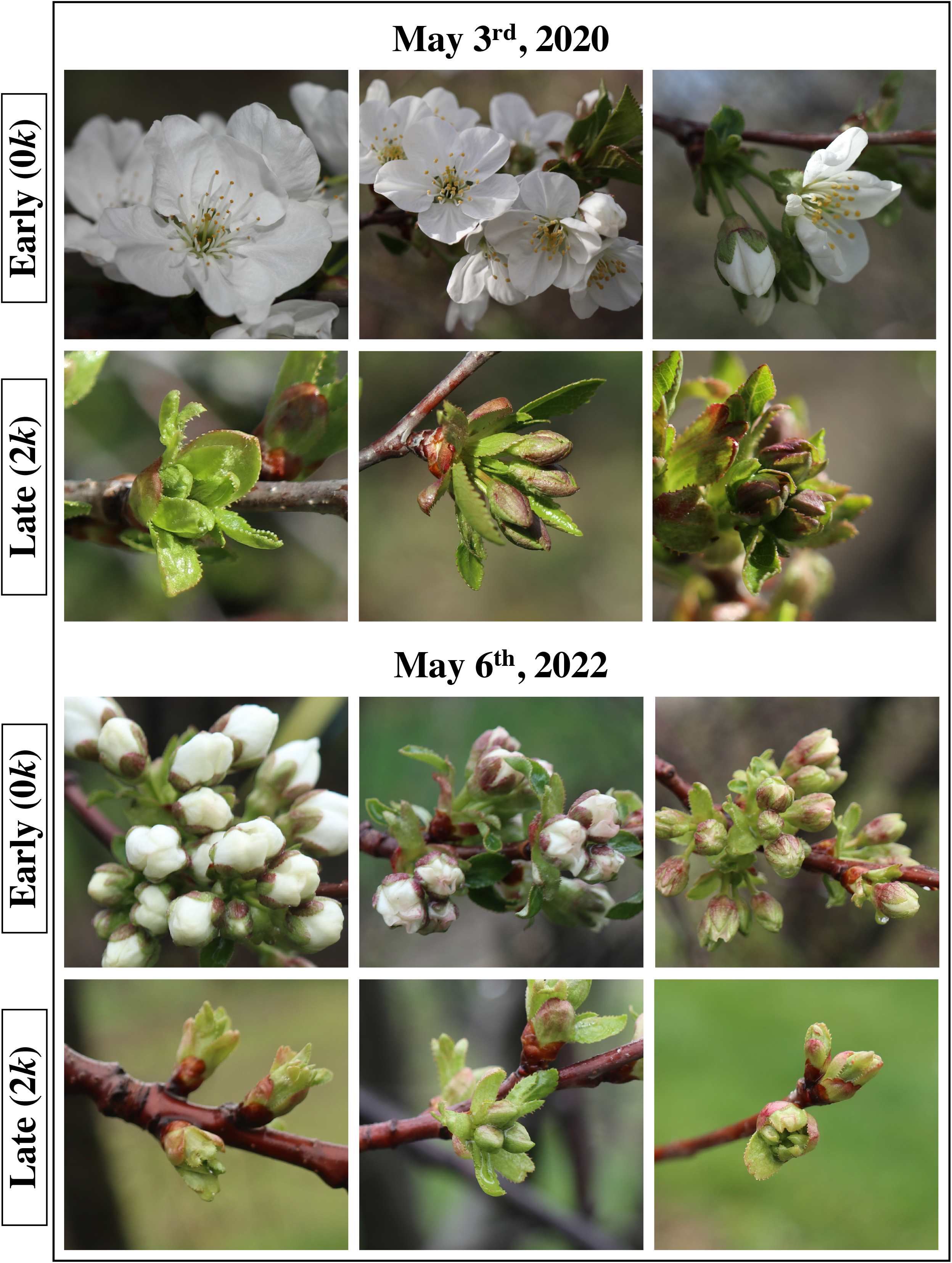
Floral bud photos of three late-blooming (inherited 2 *k* haplotypes, one from each parent) and three early-blooming individuals (inherited 0 copies of *k*) on the same day for two years. The top three individuals for each date are Early 1, Early 2, and Early 4, respectively. The bottom three individuals for each date are Late 3, Late 1, and Late 2, respectively.

A preliminary observation of segregating individuals in the late winter of 2018 revealed slight differences in internal development between early-blooming (EB) and late-blooming (LB) siblings even as the flowers were still dormant. For example, EB anthers were further advanced based on a more defined shape and differentiation of cell layers **Figure S1**. Therefore, flowers/apices in the floral position of several early-blooming and late-blooming sour cherry trees from the same population were collected from June to May of the 2018-2019 and 2019- 2020 seasons to determine how and when these developmental changes occurred. For effective comparisons between early- and late-blooming siblings, we developed a novel floral development staging system for sour cherry totaling 21 stages that span the entire life of the flower (**Figure 2**, **Figure 3**). The staging system was developed as previous works usually described only a portion of floral development or were based on external phenology (Diaz *et al*., 1981; Foster *et al*., 2003; Engin and Ünal, 2007; Mimida *et al*., 2011; Fadón *et al*., 2015, 2018*b*; Yarur *et al*., 2016; Vimont *et al*., 2019; Fernandez *et al*., 2020; Villar *et al*., 2020; Goeckeritz *et al*., 2023*a*). We anticipate our staging system will be applicable to other *Prunus* species since basic developmental patterns of individual flowers are often similar, although the structure of the inflorescence may vary (Guimond *et al*., 1998; Engin and Ünal, 2007; Fadón *et al*., 2015; Wang *et al*., 2019).

**Figure 2:**
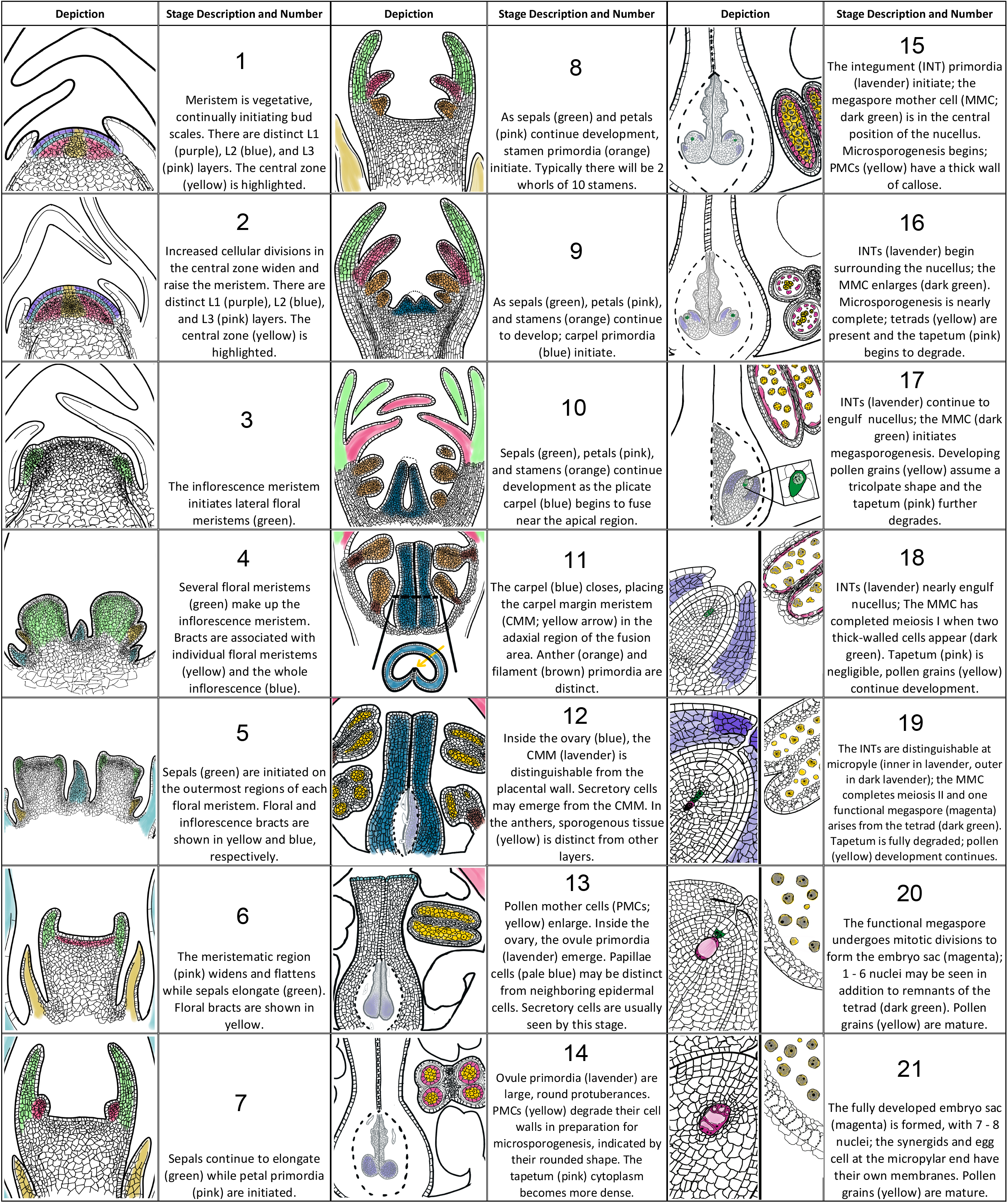
A floral development staging system for sour cherry describing development pre-floral initiation to just before anthesis. A stage number and description are given to the right of each respective depiction. Colors are used to highlight features important for stage classification.

**Figure 3:**
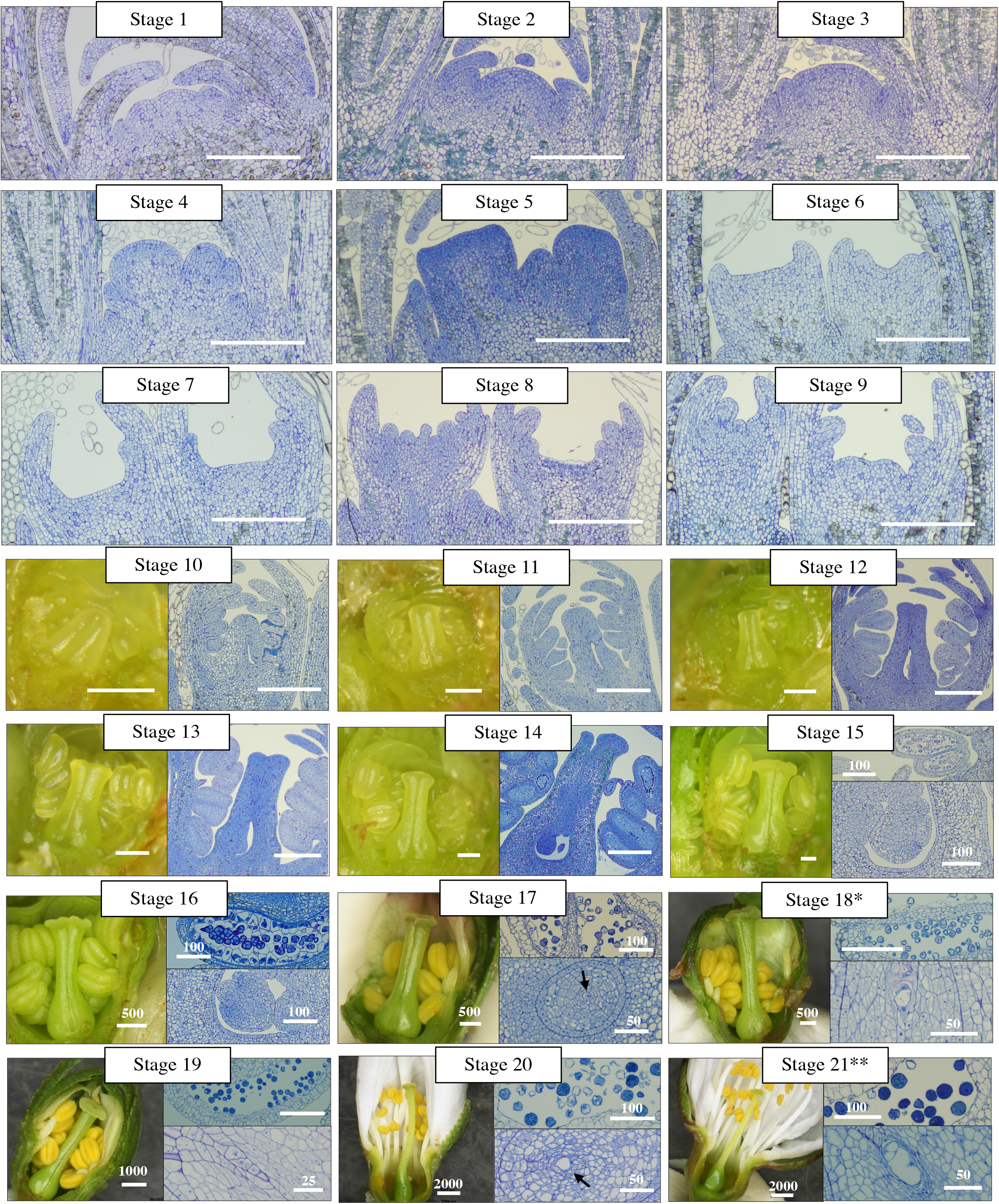
Photos of histological sections and hand dissections illustrating each stage described in Figure 2. An * indicates these section photos are identical to those in Figure 4.8E. A ** means the anther / pollen photograph is identical to that in Figure 4.8A, and the embryo sac is a different section of the same sample also in Figure 4.8A. Scale bars are 200 µm unless stated otherwise.

### Early- and late-blooming individuals differed in development as early as floral initiation

Histological comparisons between bloom groups revealed the first detectable developmental differences in EB and LB siblings of this segregating population appear as early as floral initiation and persist throughout the year. In June 2018 and 2019, the apices from both bloom groups were vegetative, continually initiating bud scale primordia (Stage 1) (**Figure 4**: 6/12/19, 6/29/19; **Figure S2**). In August 2018, apices of the different bloom groups were well into organogenesis and EB trees were more advanced than their late siblings by an average of three stages (**Figure S3**). This necessitated a finer series of collections for the summer of 2019, which revealed that on average, EB individuals committed to floral development (Stage 2+) before their LB counterparts (**Figure 4**: 7/18/19). As organogenesis progressed into dormancy these slight disparities were maintained, and the LB individuals did not surpass their EB siblings in either year (**Figure 5, Figures S3 – S6**). Still, even as relative development was consistent between the bloom groups, the same individuals observed on 8 Aug 2018 were more advanced compared to 8 Aug 2019 (Stage 5 – 9 vs Stage 4 – 7).

**Figure 4:**
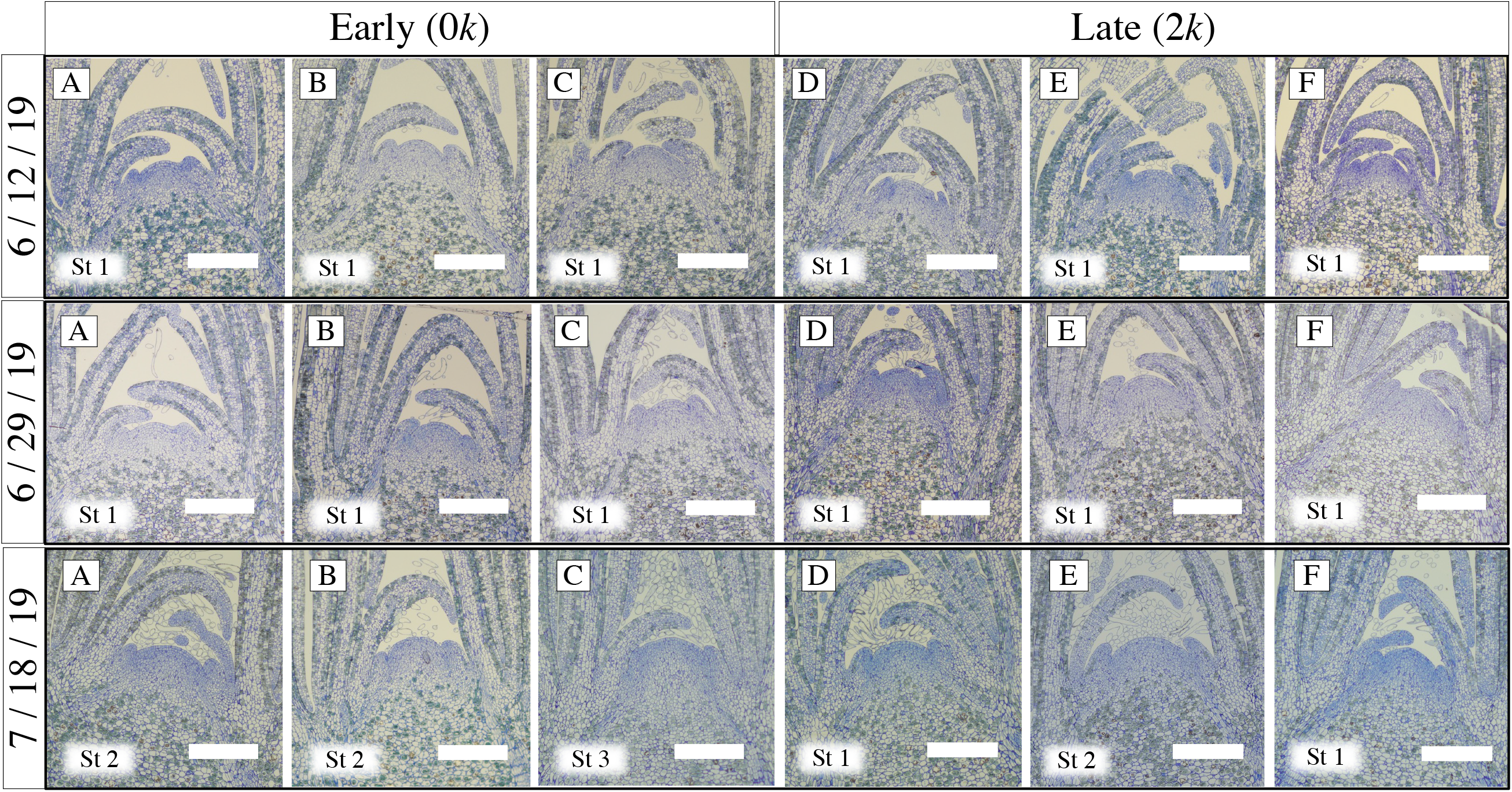
Early-bloomers are the first to make the vegetative to floral transition. Histological sections illustrate the stage of the apex from 6/12/19 – 7/18/19 based on the system in Figure 4.2. Sampling date is indicated on the y-axis. Individuals A – C are Early 1, Early 2, and Early 3. Individuals D – F are Late 1, Late 2, and Late 3. All apices appeared morphologically vegetative until 18 July, when more early-blooming apices on average made the floral transition. These three dates and individuals of each bloom group were used for the RNA-sequencing analyses. Scale bars are 200 µm unless stated otherwise.

**Figure 5:**
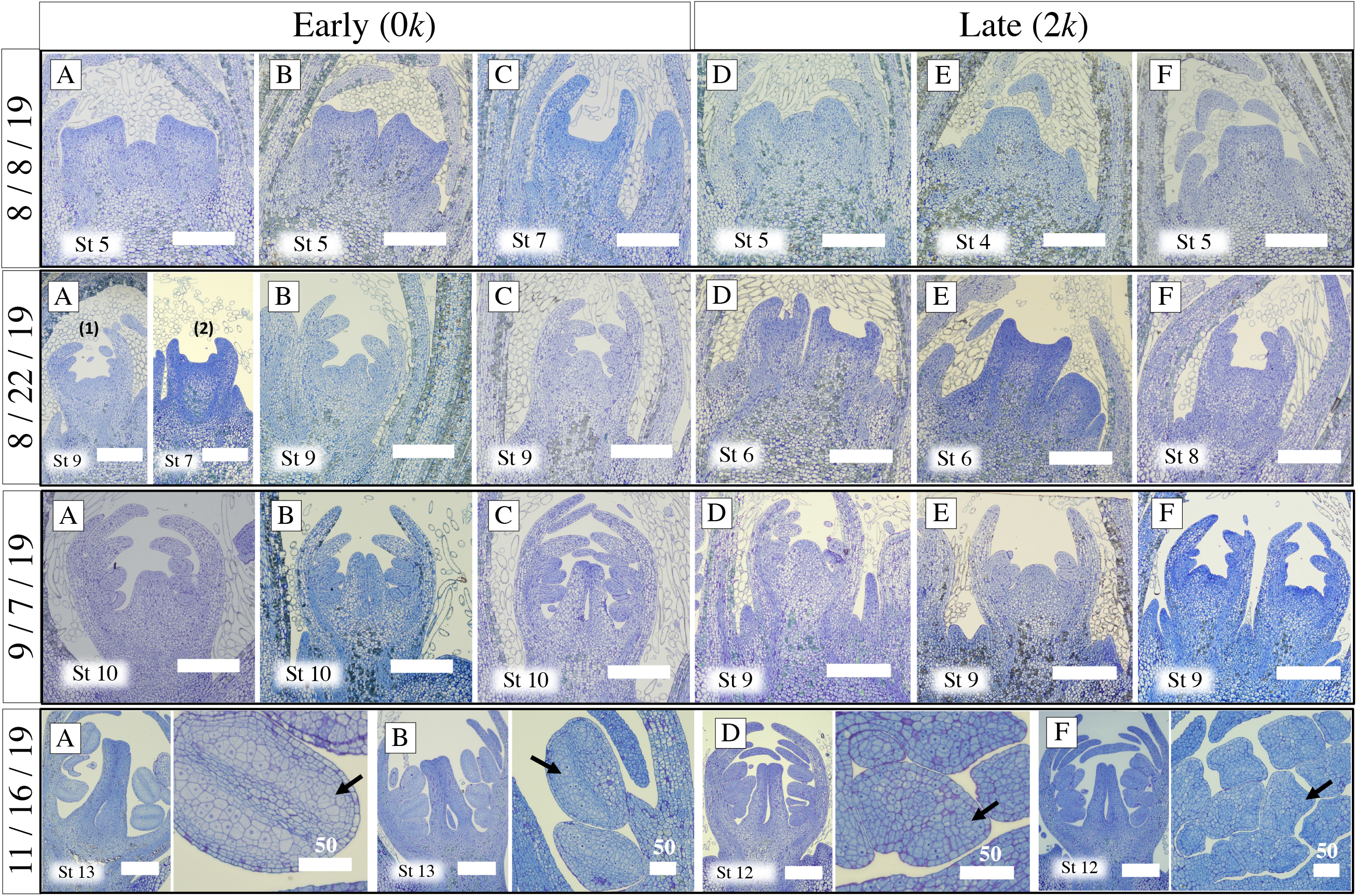
Early-bloomers are more advanced as floral organogenesis proceeds to dormancy. Histological sections illustrating the stage of floral development from 8/8/19 – 11/16/19 based on the system in Figure 2. Sampling date is indicated on the y-axis. Individuals A – C are Early 1, Early 2, and Early 3, respectively. Individuals D – F are Late 1, Late 2, and Late 3, respectively. Note that only four individuals were sectioned for 11/16/19. Early 1 (A) on 8/22/19 is represented by two photos to show the variability in this individual. Black arrows in 11/16/19 indicate sporogenous tissue, which is more defined in the early-bloomers. On all of these dates, early-bloomers keep their lead in development. Scale bars are 200 µm unless stated otherwise.

As the flowers entered dormancy, EB trees were at similar stages in both years (Stages 12 – 13; **Figure 5**: 11/16/19; **Figures S6 – S7**). Sporogenous tissue was easily discernible from the other anther layers, and the protuberance from the placental wall marked the carpel margin meristem (CMM; Stage 12) and the start of ovule formation (Stage 13). Late-bloomers were also at similar stages between the two years (Stages 11 – 12), but with sporogenous tissue barely distinguishable from other anther layers and a rudimentary CMM at best (**Figure 5**: 11/16/19; **Figures S6 – S7**). Winter proceeded with little development until flowers were close to exiting dormancy (**Figure 6**; **Figures S8 – S9**). At this time, increasing amounts of elongated cells emerged from the CMM sometime between carpel closing and the appearance of stylar tracts (**Figure 6B, E, H**; **Figure S8B, C; Figure S9A – C**). Given the timing of their appearance, it is speculated they may be involved in the development of stylar tissue.

**Figure 6:**
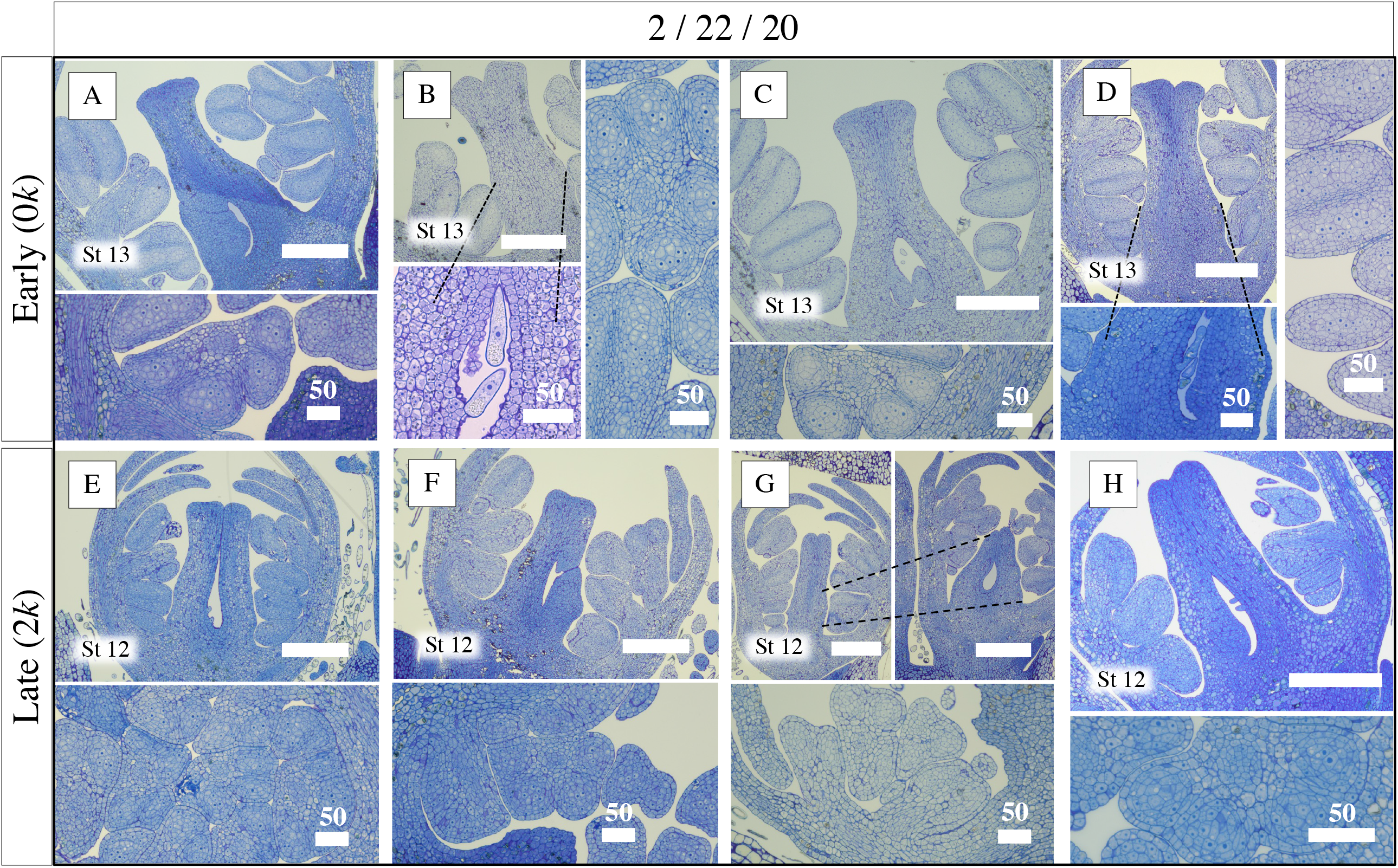
Early-bloomers remain ahead in their floral development when both bloom groups exit dormancy. Histological sections illustrate developmental stages on 2/22/2020, just before warm temperatures return. A – D represent Early 1, Early 2, Early 4, and Early 5, respectively, while E – H represent Late 1, Late 2, Late 3, and Late 4, respectively. Dotted lines indicate separate sections of the same flower to show the internal state of the ovary. The cell layers within the early-blooming anthers are more differentiated compared to those in late-blooming flowers, although sporogenous tissue is at least visible for all genotypes. The ovule is a noticeable protrusion in early-bloomers while most late-bloomers’ flowers have rudimentary carpel margin meristems. Scale bars are 200 µm unless stated otherwise.

Growth resumed exponentially in the spring and flowers underwent microsporogenesis before megasporogenesis (**Figures 7 – 8; Figures S10 – S11**). As seen at other time points, the EB trees persisted ahead of their LB siblings and bloomed first. On 6 Apr 2020, EB individuals continued pollen development and the megaspore mother cell became distinct from the surrounding tissues in the nucellus (Stages 15 – 17; **Figure 7A – D**). Meanwhile, LBs were just initiating microsporogenesis and integument primordia were rarely seen (**Figure 7E – H**). Occasionally, the stage of microsporocyte development was slightly out of sync compared to other features of the flower (**Figure 7A, D**; **Figure 8G**). For EBs on 2 May 2020, pollen was mature, the tapetum was fully degraded, and the megagametophyte was in the process of forming the embryo sac (Stages 20 – 21; **Figure 8A – D**). However, remnants of the tapetum endured in the majority of LBs; and in all LBs, the megasporophyte ranged from just beginning megasporogenesis to just completing it (Stages 17 – 19; **Figure 8E – H**).

**Figure 7:**
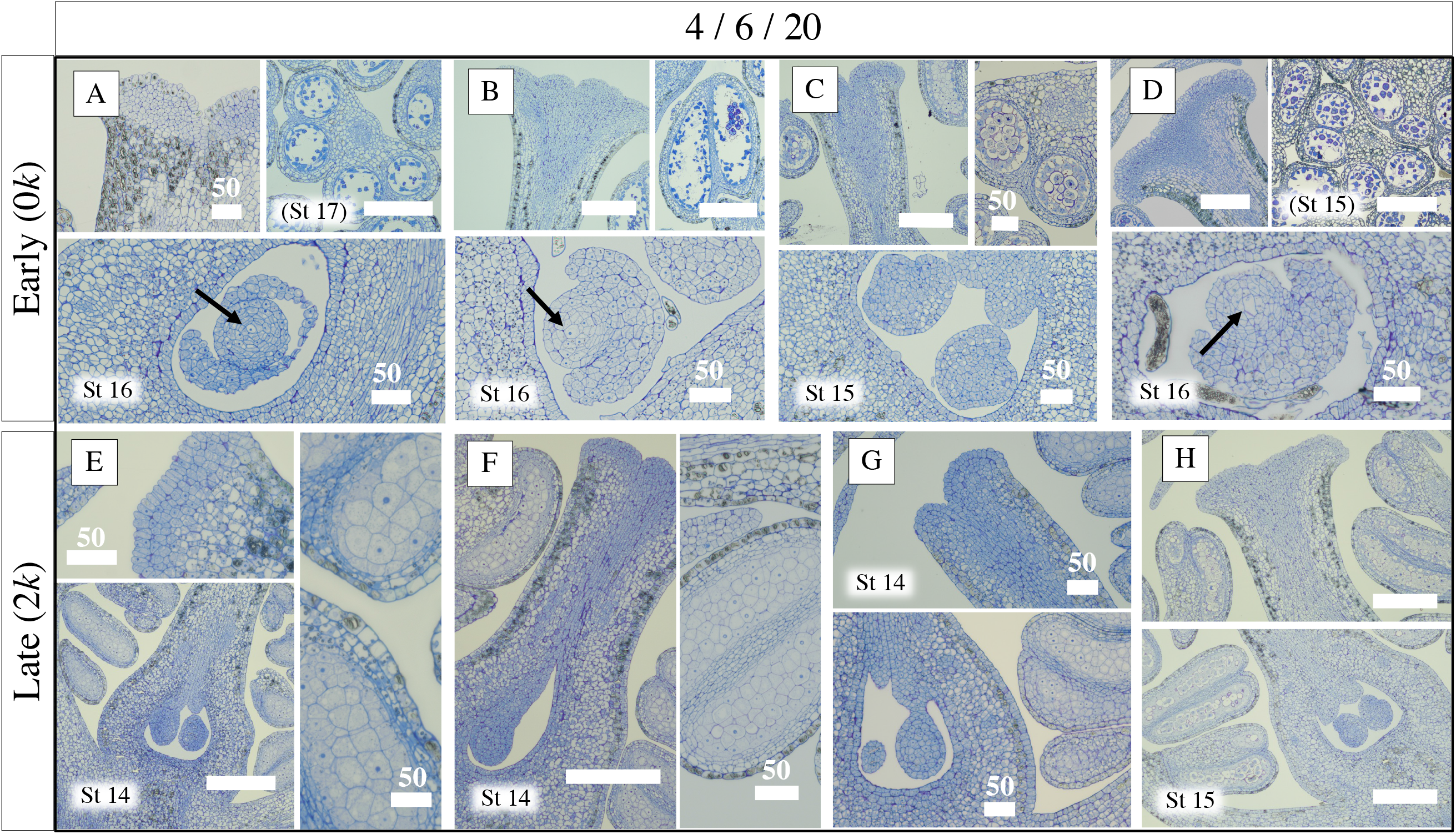
Early-blooming and late-blooming flowers are at different stages of ovule formation and microsporogenesis in early spring. Histological sections illustrating developmental stages on 4/6/2020. A – D represent Early 1, Early 2, Early 4, and Early 5, respectively, while E – H represent Late 1, Late 2, Late 3, and Late 4, respectively. Arrows indicate the emerging megaspore mother cell at the center of the nucellus for several early-bloomers; this cell is not yet visible in late-bloomer flowers. Most late-bloomer flowers have not undergone microsporogenesis while most early-bloomer flowers have. Scale bars are 200 µm unless stated otherwise.

**Figure 8:**
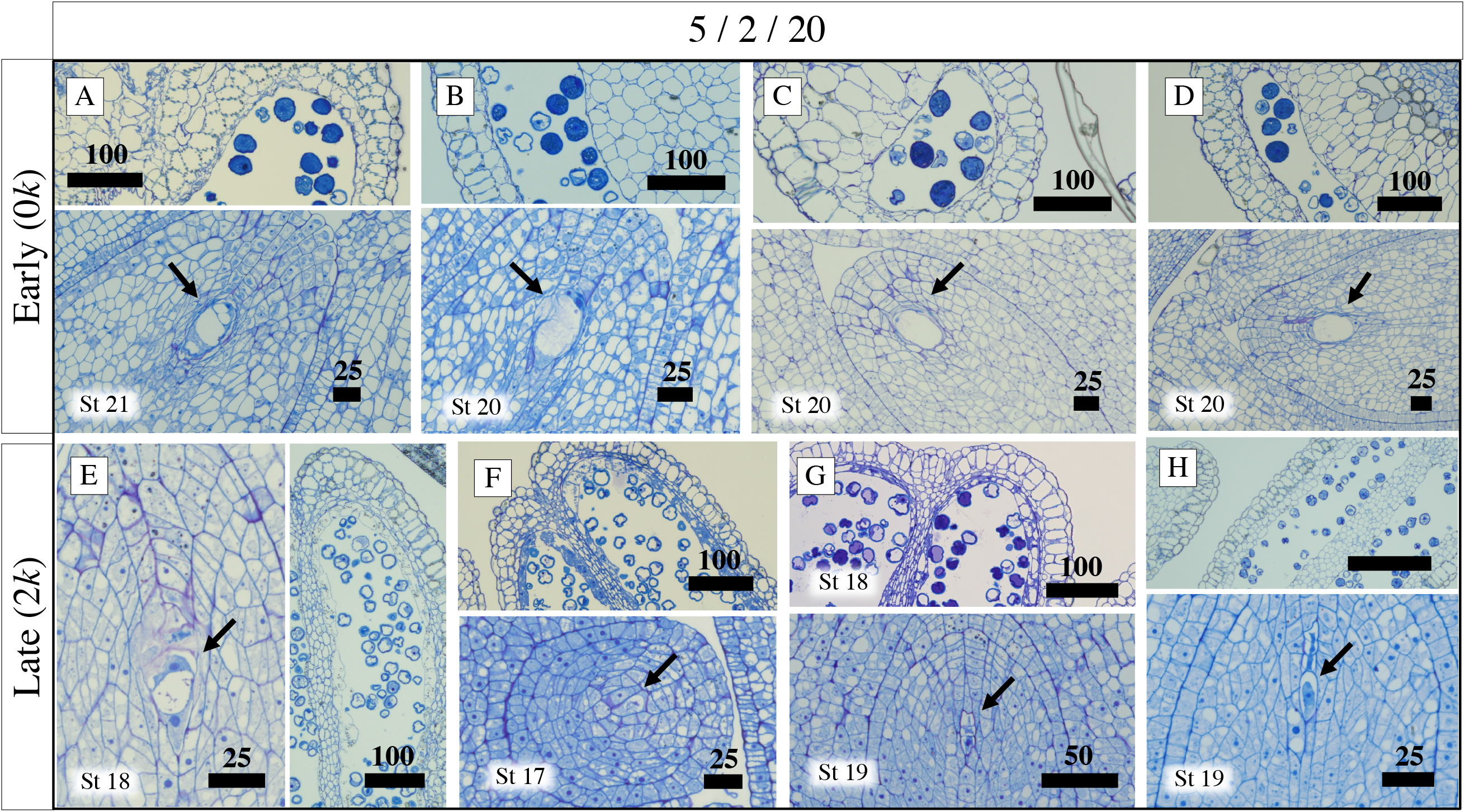
Early-blooming and late-blooming individuals are at different stages of megasporogenesis in mid-spring. Histological sections illustrating developmental stages on 5/2/2020, the day before full bloom for Early 1. A – D represent Early 1, Early 2, Early 4, and Early 5, respectively, while E – H represent Late 1, Late 2, Late 3, and Late 4, respectively. Arrows indicate the megagametophyte, which ranges in development from just initiating meiosis (late-bloomer) to a fully-developed embryo sac (early-bloomer). Scale bars are 200 µm unless stated otherwise.

Lastly, to discern the significance of the qualitative differences observed between EBs and LBs during the period of floral initiation and the onset of dormancy, we carried out two-sample Wilcoxon rank sum tests on nine dates: 8 Aug 2018, 20 Sep 2018, 30 Oct 2018, 28 Nov 2018, 18 Jul 2019, 8 Aug 2019, 22 Aug 2019, 7 Sep 2019, and 16 Nov 2019. Statistical significance (Bonferroni corrected p-value < 0.05) was detected between the EB and LB stage designations at the pre-dormancy dates of 20 Sep 2018, 30 Oct 2018, 8 Aug 2019, 22 Aug 2019, and 7 Sep 2019. However, upon entering dormancy (28 Nov 2018 and 16 Nov 2019), differences in developmental stage were not statistically significant. A graphical summary of the histological results is presented in **Figure 9** for the seasons of 2018-2019 and 2019-2020.

**Figure 9:**
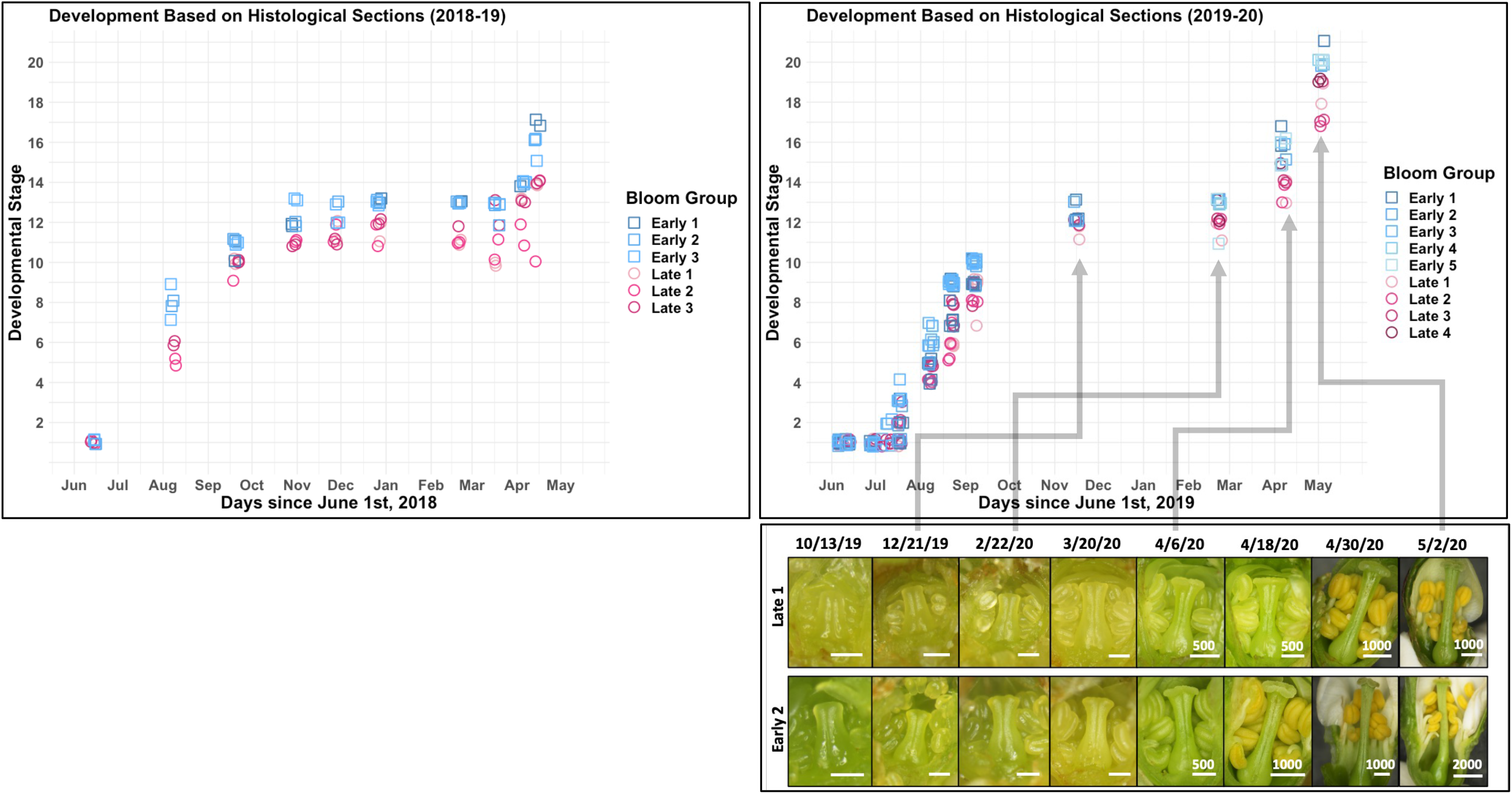
A summary of the histological results for early- and late-blooming siblings for two years show differences as early as floral initiation that persist until bloom. Qualitative developmental stage (see Figure 2) is plotted against calendar date. The lull in each sigmoidal curve represents dormancy, when very little visible development occurred and very few GDDs accumulated. Lastly, dissection photos in the bottom right illustrate floral development of a late-bloomer (top row) and early-bloomer (bottom row) on select dates; some of these dates are indicated on the graph above. Early-blooming siblings, those that inherited no k haplotypes, begin developmental divergence around the time of floral initiation and maintain this lead for the rest of the year. Scale bars are 200 µm unless stated otherwise. A ‘jitter’ function was used in R for clearer visualization of data points.

**Figure 10:**
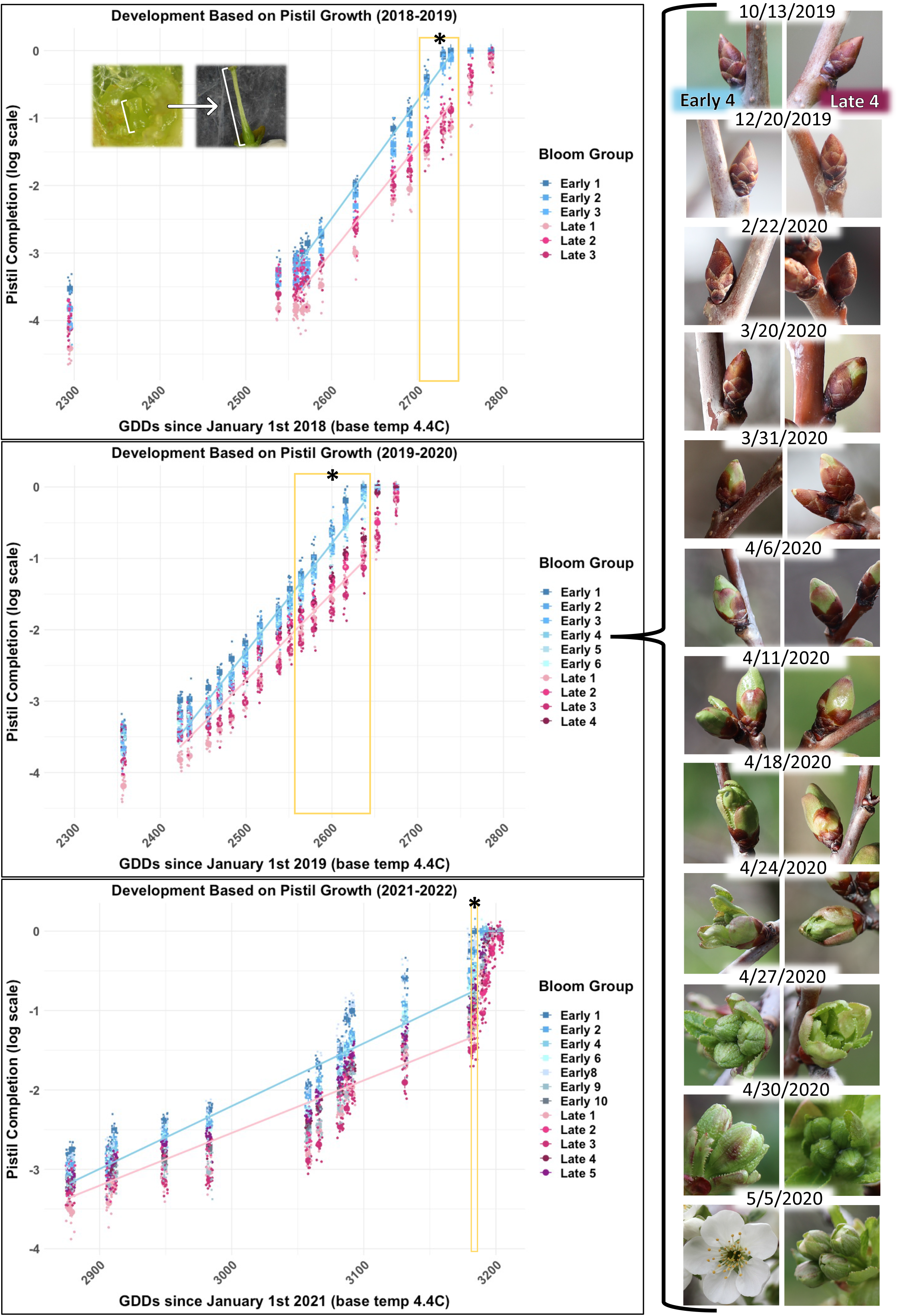
Pistil measurements of early- and late-blooming siblings for three years support small pre-dormancy differences and variation in spring growth rate. Pistil measurements are indicated on the left, with growing degree days (GDDs; base temperature 4.4 °C) since January 1st of the year of floral initiation plotted on the x-axis and pistil completion (log scale) on the y-axis. Trees are colored by bloom group (blues = EB; pinks = LB) with means plotted as large symbols. Smaller symbols indicate distinct measurements for the respective individual. A ‘jitter’ function was used in R for clearer visualization of data points. Slopes of growth trends for each bloom group are approximated from dormancy to the date the first tree had reached bloom. Trees reaching 0 on the log scale first in less GDDs bloom earlier. The inset in the top left demonstrates how pistils were measured. Measurements were done from 9/20/18 – 5/16/19, 10/13/19 – 5/15/20, and 12/29/21 – 5/13/22. Gold boxes with asterisks indicate collection dates / GDD values where a statistically significant difference between the bloom groups was detected (Type III ANOVA; p < 0.05). In all years, GDDs, bloom group, and their interaction significantly impacted developmental rate. Notice the different x-axis scale for 2021 – 2022 – more GDDs were accumulated in this season. The photos on the right are representatives of Early 4 and Late 4’s floral buds on select dates of the 2019 – 2020 season.

### Pistil progression supports consistent differences between EB and LB individuals

To quantify and compare development between EB and LB siblings, pistil growth was measured for the 2018-2019, 2019-2020, and 2021-2022 seasons (**Files S1 – S3**). Since the pistil primordia do not emerge until late summer or fall, this limited possible measurements to this developmental window.

Despite the variability in sample size between years, the pattern of development was consistent (**Figure 10**). In all three years, small differences between bloom group prior to dormancy were detected, although they were statistically insignificant. Once growth resumed in the spring with the accumulation of GDDs, the rate of development was significantly faster for the EBs compared to the LBs. Boxed dates in **Figure 10** emphasize when statistical significance was detected at the p < 0.05 level between the bloom groups according to a Tukey’s HSD test. Individuals reaching 0 on the log scale (representing an open flower) with fewer GDDs bloomed earlier. Significance between the bloom groups was detected between 2711 – 2739 GDDs in 2019, 2564 – 2637 GDDs in 2020, and only at 3184 GDDs in 2022. The corresponding calendar dates for the three years were 4 May – 8 May, 24 Apr – 5 May, and 8 May, respectively. The reason for the shrunken range in bloom times and the spike in development across fewer collections in late spring of 2022 (∼3100 – 3200 GDDs) was likely due to a string of unseasonably warm days that accelerated development.

The quantitative pistil measurements are consistent with histological observations since they imply variation between bloom groups was established prior to dormancy onset for all years. These differences apparently became more pronounced in spring when the growth rate was exponential.

### Budbreak forcing experiments suggest contrasting temperature perception between early- and late-blooming siblings

Classical forcing experiments were performed to assess the depth of dormancy and compare temperature perception of early- and late-blooming siblings. Dormant branch cuttings with floral buds were collected from individuals in each bloom group periodically for two winters (**Table S4**). Each collection represented a certain amount of chill floral buds had sensed up to the date of collection. Budbreak on cuttings was recorded as the percent of flower buds reaching each stage on the BBCH scale, which is a decimal coding system for the phenology of woody perennials (Fadón *et al*., 2015). The BBCH system was used since it is simple, non-destructive, and does not require flower dissections. Trends presented in **Figure 11** demonstrate the bloom groups’ responses given a certain level of chill (x-axis) and a constant, average amount of GDDs across time points for each year (121 GDDs for 2019 – 2020, 96 GDDs for 2021 – 2022).

**Figure 11:**
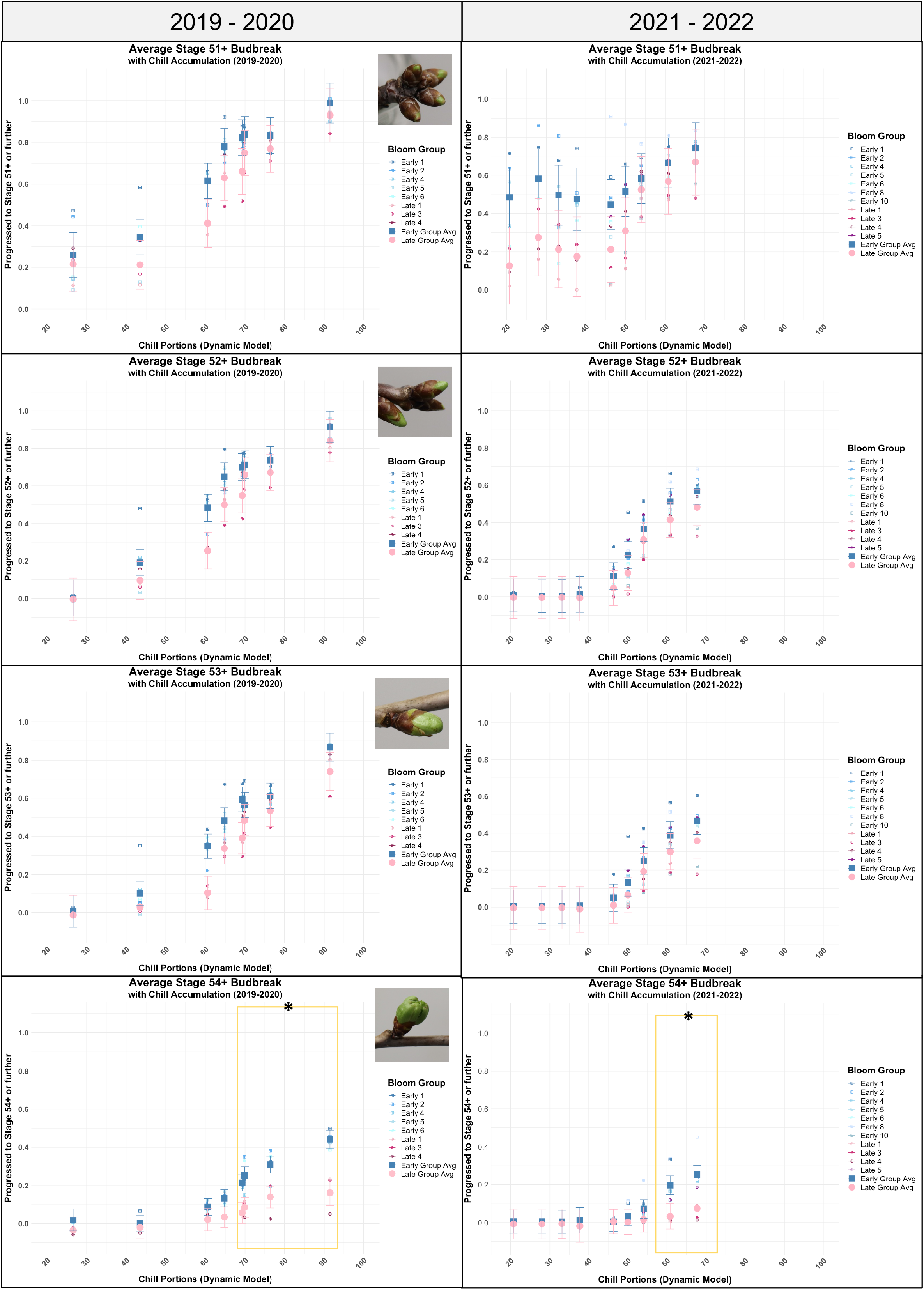
Early-bloomers respond first to warm temperatures and additional chill equates to a faster rate of growth compared to the late-bloomers. Forcing experiments were conducted for two winters. The GDDs for each response are a constant average within years (121 GDDs for 2019 – 2020; 96 GDDs for 2021 – 2022) Each column represents the winter of one year, and rows represent flower percentages reaching stages 51, 52, 53, and 54. Stages were modeled separately to show the gradual budbreak responses of the bloom groups given the same amount of chill and forcing conditions. Gold boxes with asterisks indicate statistical differences between the bloom groups when the means were separated by chill level. Axes are consistent for visualization of the trends between years. Means for each bloom group are shown as solid blue squares or pink circles and the error bars are the 95% confidence intervals of the statistical model. Smaller symbols represent the means for each tree. Inset photos in the upper right corners of the 2019-2020 data are a representative photo for the respective BBCH stage. Note that since 2019 – 2020’s average GDD accumulation was slightly higher, this likely explains the increase in budbreak for all chill levels and stages compared to 2021 – 2022 (i.e., an average of 121 GDDs vs 96 GDDs).

BBCH Stage 51 is characterized by swelling of the dormant bud and the first sign of green tissues beneath the bud scales. In 2019 – 2020, the main effects of bloom group, chill, and GDDs had a significant effect on whether flowers progressed to stage 51, with EB individuals having an average of 11% more flowers progressing to this stage compared to the LB trees (p < 0.05, Type III ANOVA and Tukey’s HSD; **Figure 11**). The interaction of GDDs and chill was also statistically significant. In 2021 – 2022, the main effects of bloom group, GDDs, their interaction, and the interaction between GDDs and chill were also significant. However, the main effect of chill was not (p = 0.30). After a given amount of heat, EBs averaged 21% more flowers reaching stage 51 compared to LBs. The average differences seen in both years are reflected in the growth trends: when flowers begin responding to forcing temperatures, the mean budbreak percentage for the early-bloomers is always slightly higher (**Figure 11**). This observation suggests EBs tend to respond to forcing before their LB siblings irrespective of how much chilling has accumulated.

BBCH Stages 52 and 53 are defined by further bud swelling and exposure of green tissues. BBCH 52 was not a formal stage prior to the current study, so it was created here as an intermediate between 51 and 53 to add resolution between stages. In both years, the main effects of GDDs and their interaction with chill were major developmental drivers to these intermediate stages. However, the main effect of bloom group only made the significance cutoff in 2019 – 2020 for BBCH 52. For stage 53 of this year and both stages in 2021 – 2022, bloom group’s interaction with GDDs was statistically significant in driving budbreak. Chill was significant at the p < 0.05 level only in 2019 – 2020 as well. The reason for the discrepancies among the two years for the main effects of chill and bloom group may be due to increased variability as more individuals were added in the 2021 – 2022 season. Moreover, the design was less balanced since a few individuals were not sampled at every date (chill level; see **Table S4**). However, one can conclude the response trends agree for both stages and years (**Figure 11**).

Lastly, BBCH Stage 54 is ‘budbreak’ in the most literal sense as it marks the first opening of the flower bud. In both seasons, the same effectors were statistically significant in influencing development to this stage: the main effects of bloom group and GDDs, the interaction of bloom group and GDDs, the interaction of bloom group and chill, and the interaction of chill and GDDs. Effectively what this means is that given a certain amount of chilling, forcing translates to a faster rate of development for early-bloomers. Interestingly, these forcing results echo those of the pistil growth analyses. However, the analyses differ in their sensitivities: the more quantitative assessment of pistil growth suggested minor differences prior to dormancy, but development according to the BBCH scale placed the EB and LB groups at the same stage. This stresses the consequences of scale when it comes to developmental comparisons and their subsequent conclusions. Still, in traditional terms, one may conclude the EBs have lower chilling requirements than their LB siblings, and they perceive temperature differently.

### Higher quantities of ROS associate with depth of dormancy irrespective of bloom group

Reactive oxygen species (ROS) have been proposed as a potential growth regulator for dormancy transitions, with higher ROS content being associated with deeper dormancy (Javier *et al*., 2021). Therefore, we were interested in whether the segregating individuals differed in their ROS content during dormancy. Hydrogen peroxide (H_2_O_2_), a relatively stable ROS, was quantified in floral buds collected during the late fall and winter of 2019 – 2020 between 13 Oct 2019 (6.6 CP since 1 Oct 2019) and 23 Mar 2020 (93.4 CP) for five dates (**Table S5, File S6**). According to a type III ANOVA, bloom group did not significantly influence H_2_O_2_ concentrations (p = 0.15); however, chilling (synonymous with collection date) did (p = 4.9 x 10^-5^; **Figure 12)**. µmoles of H_2_O_2_ per gram of fresh weight were similarly lower on 13 Oct, 22 Feb, and 23 Mar compared to 16 Nov and 20 Dec. The higher concentrations of H_2_O_2_ coincided with deeper dormancy, as determined by the response in forcing conditions. The lower concentrations appeared to be linked with visible growth and / or with the speed at which flower buds respond to forcing temperatures. It should be noted that H_2_O_2_ showed a weak negative correlation with temperature at the time of collection (**Figure S12**). However, this association should be interpreted with caution since this is to be expected if chill accumulation is equivalent to the time of year. Moreover, the range of temperatures during our collections was narrow; therefore, temperature on the day of collection was excluded from the statistical model as a more thorough assessment is needed to detect any sort of relationship. Nonetheless, these observations support the hypothesis that higher ROS content reinforces dormancy and is inhibitory to growth.

**Figure 12:**
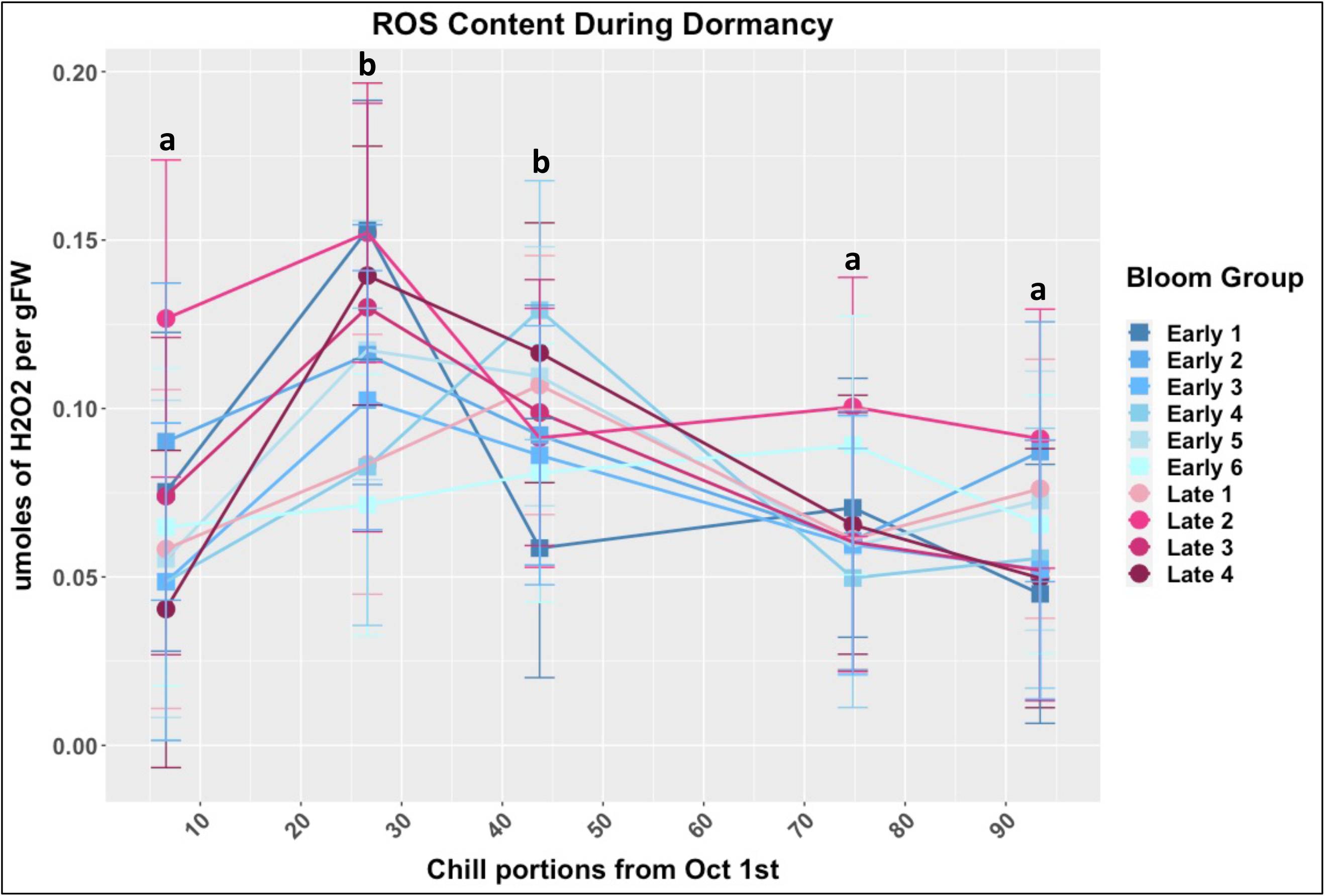
Concentration of hydrogen peroxide (H_2_O_2_), a reactive oxygen species, associates with depth of dormancy. H_2_O_2_ content (y-axis) was measured five times from fall (13 Oct) to spring (23 Mar) to determine if there were differences between the bloom groups as they accumulated chill (x-axis) and navigated dormancy. While bloom group did not significantly affect H_2_O_2_ content, chill levels (collection date) did. The significantly higher concentrations (signified with ‘b’) represent early winter collections when flowers hardly responded to forcing conditions, (see forcing experiment results for 2019 – 2020). Means for each genotype are shown as blue squares or pink circles and the error bars are the 95% confidence intervals of the statistical model.

**Figure 13:**
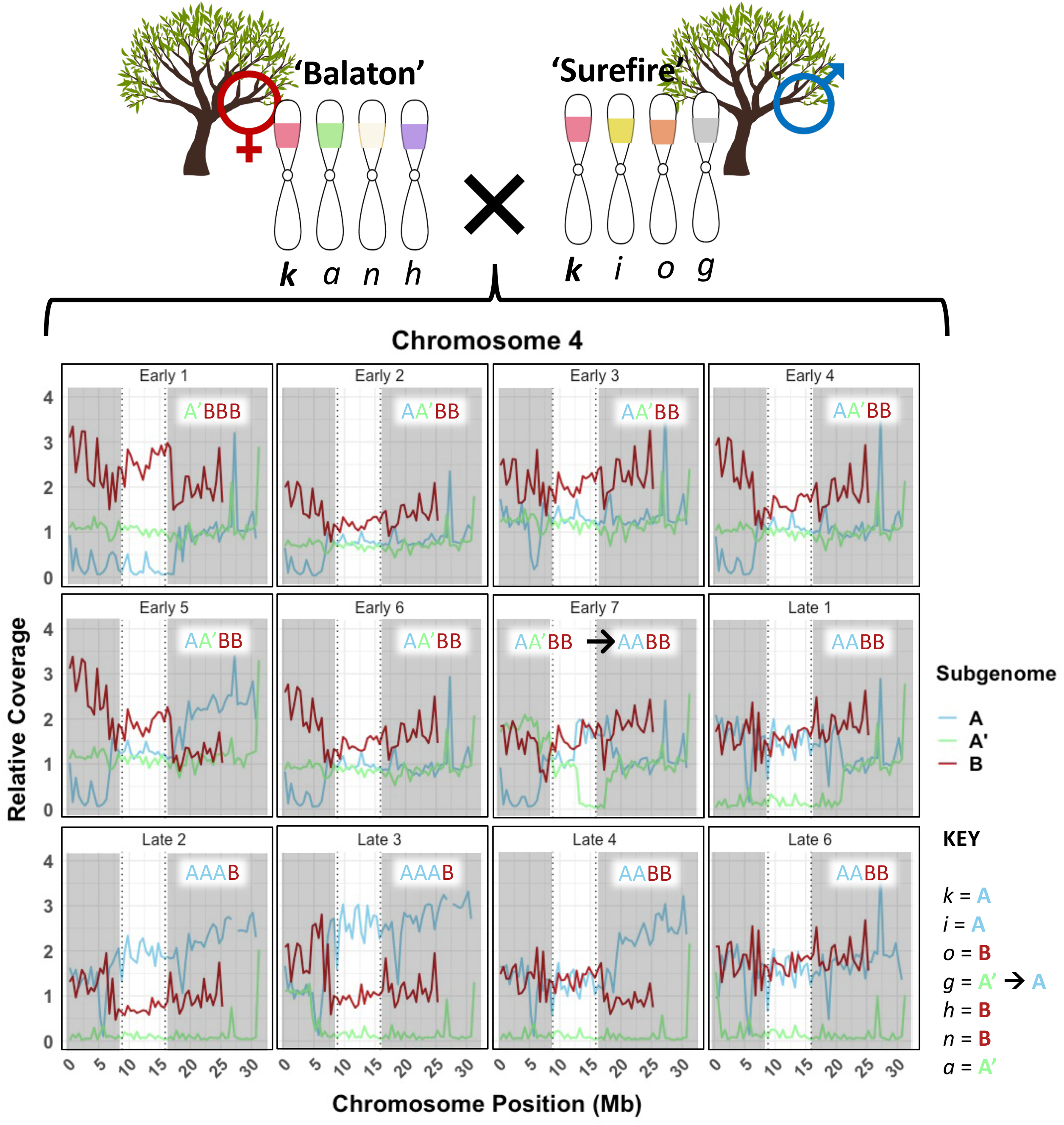
Comparing coverage of each subgenome at the bloom time QTL suggests the late-blooming haplotype *k* originates from subgenome A. Subgenome dosage at the bloom time QTL was determined by mapping gDNA Illumina reads of segregating individuals and their parents to the ‘Montmorency’ reference genome. Using haplotype information from Cai et al. 2018, pairwise comparisons of dosage and haplotype composition revealed the subgenome each haplotype most likely originates from (**File S7**). The bloom time QTL for the 12 siblings’ plots is left unshaded and separated by the rest of the chromosome with dotted lines. Subgenome dosages are given at the top right corner or top center (Early 7) of each plot.

### A genomic coverage analysis of EB and LB individuals suggests k is derived from Prunus cerasus subgenome A

Recent findings implied different accessions of sour cherry contain various dosages of the three subgenomes (A, A’, and B) due to homoeologous exchanges and/or subgenome chromosome replacements (personal communication with Iezzoni and Rhoades). Thus, we reasoned that the parents and individuals of the segregating population could have different dosages at the QTL, and exploring the subgenome content of this region may yield insight into which subgenome(s) the late-bloom *k* haplotype is derived from. Through pairwise comparisons and deductive reasoning (fully detailed in **File S7**), *k* was determined to originate from subgenome A, a subgenome donated by the late-blooming *Prunus fruticosa-*like progenitor. For example, according to the relative coverage analysis, Early 1 has dosage A’BBB at the QTL (**Figure 13**). The previous mapping study (Cai *et al*., 2018) showed this individual to have a haplotype composition of (*a, h, o, o*). Because B is the only subgenome with at least 2X dosage, haplotype *o* must come from subgenome B. Furthermore, since there is no subgenome A dosage in the QTL for this genotype, haplotypes *a* and *h* do not originate from subgenome A (**File S7**). Notably, the sequenced LB siblings showed no dosage of A’, meaning they did not inherit haplotypes *a* or *g*. This might partially be explained by the preferential pairing of subgenomes during meiosis, a well-documented occurrence in sour cherry (Beaver and Iezzoni, 1993; Tsukamoto *et al*., 2010).

### Genetic variation relevant to the k haplotype

After aligning gDNA reads of the individuals and parents to the three ‘Montmorency’ subgenomes, single nucleotide polymorphism (SNPs) and small insertions and deletions (INDELs < 30 bp) were called in an ∼8 Mb region encompassing the QTL – only calling variants in individuals with some dosage of subgenome A. Ultimately this meant comparisons were between haplotypes *k*, *i*, and the latter part of *g* since the other haplotypes originated from subgenomes A’ and B. After hard-filtering for quality of the variant calls, the dataset was further filtered based on *k*’s uniqueness from the other subgenome A haplotypes. This analysis resulted in 8101 variants. According to snpEff, 7726 of these variants were classified as ‘modifiers,’ which contained upstream, downstream, 5’ and 3’ UTR, intragenic, intergenic, and intronic variants. 128 variants were predicted to have low effects, such as start codons gained and synonymous mutations. 222 were considered moderate effect variants, which included nonsynonymous mutations and codon insertions and deletions. Finally, there were 25 high effect variants, which included those predicted to cause a new splice site, stop codon, or frameshift in the protein product. Altogether, these variants were within or associated with 755 genes. The complete list of these smaller variants is provided in **File S8**, which includes a separate tab for curated high effect variants (confirmed in reads viewed in a genome browser).

We also attempted to identify larger structural variants (SVs; > 30bp) associated with the *k* haplotype. Reads mapping to subgenome A from individuals only containing haplotype *i* or haplotype *k* were pooled to increase calling accuracy. ‘Surefire’ and Early 7, both containing haplotype *g*, were used to rule out SVs whenever possible, but relatively low coverage limited their utility (see Methods). Variants were kept even if the *k* allele was homozygous and the other allele information was missing. Post-filtering, 169 variants were found, ranging in size from 31 to nearly 81000 bp (**File S8**). Due to their length, these collective variants were predicted to affect 590 genes – the majority of genes within the QTL. This included intergenic effects, which was predicted to affect 231 genes alone. To focus time and resources, only structural variants with effects predicted to be high effect or upstream of protein-coding genes were further considered. 98 of these variants were posited to have at least one upstream effect and 18 were predicted to have high effects – many variants were speculated to affect multiple genes. Altogether, 124 genes were anticipated to have upstream SV effects, 28 were predicted to have high effect SVs, and 25 genes were predicted to be affected by both.

### Differentially expressed genes between EB and LB siblings

To narrow the list of candidate genes for the late-blooming phenotype of haplotype *k*, gene expression differences between early- and late-blooming groups around the time of floral meristem initiation were investigated. RNA from three EB and three LB individuals’ apices was sequenced on 12 Jun 2019, 29 Jun 2019, and 18 Jul 2019. Early 1, Early 2, and Early 3 constituted the EB group while Late 1, Late 2, and Late 3 constituted the LB group. Based on the haplotype key shown in **Figure 13**, the QTL subgenome dosages for the EB group trees were A’BBB, AA’BB, and AA’BB, while dosages of the LB group trees were AABB, AAAB, and AAAB, respectively, within the QTL. *Prunus cerasus* subgenome A was used as a reference for alignment given the likelihood *k* is a unique allele derived from this subgenome, and parameters were relaxed so that overall alignment rates to subgenome A were similar among individuals and time points (see Methods). We were also confident gene expression quantification biases would be minimized since *Prunus* genomes are extremely syntenic (Dirlewanger *et al*., 2004). For the first two time points, the apex in the floral position appeared vegetative (stage 1); however, on 18 Jul 2019, more apices from the EB individuals began to proceed to stage 2 and 3 (**Figure 4**).

A principal component analysis (PCA) revealed a clear separation between bloom groups, demonstrating these individuals were already distinct at the molecular level on 12 June and 29 June despite their similar morphology (**Figure 14**; **Figure 4**). Markedly, Early 1 (A’BBB) was substantially removed from its fellow EBs – perhaps suggesting unique aspects of subgenome A, especially at the chr4 bloom time locus, drive developmental changes within these tissues. While the results here concentrate on comparisons between the bloom groups, DEGs and variation in associated GO terms for comparisons among dates within bloom groups were also observed, consistent with the changes seen along PC1 (**Files S9 – S10**).

**Figure 14:**
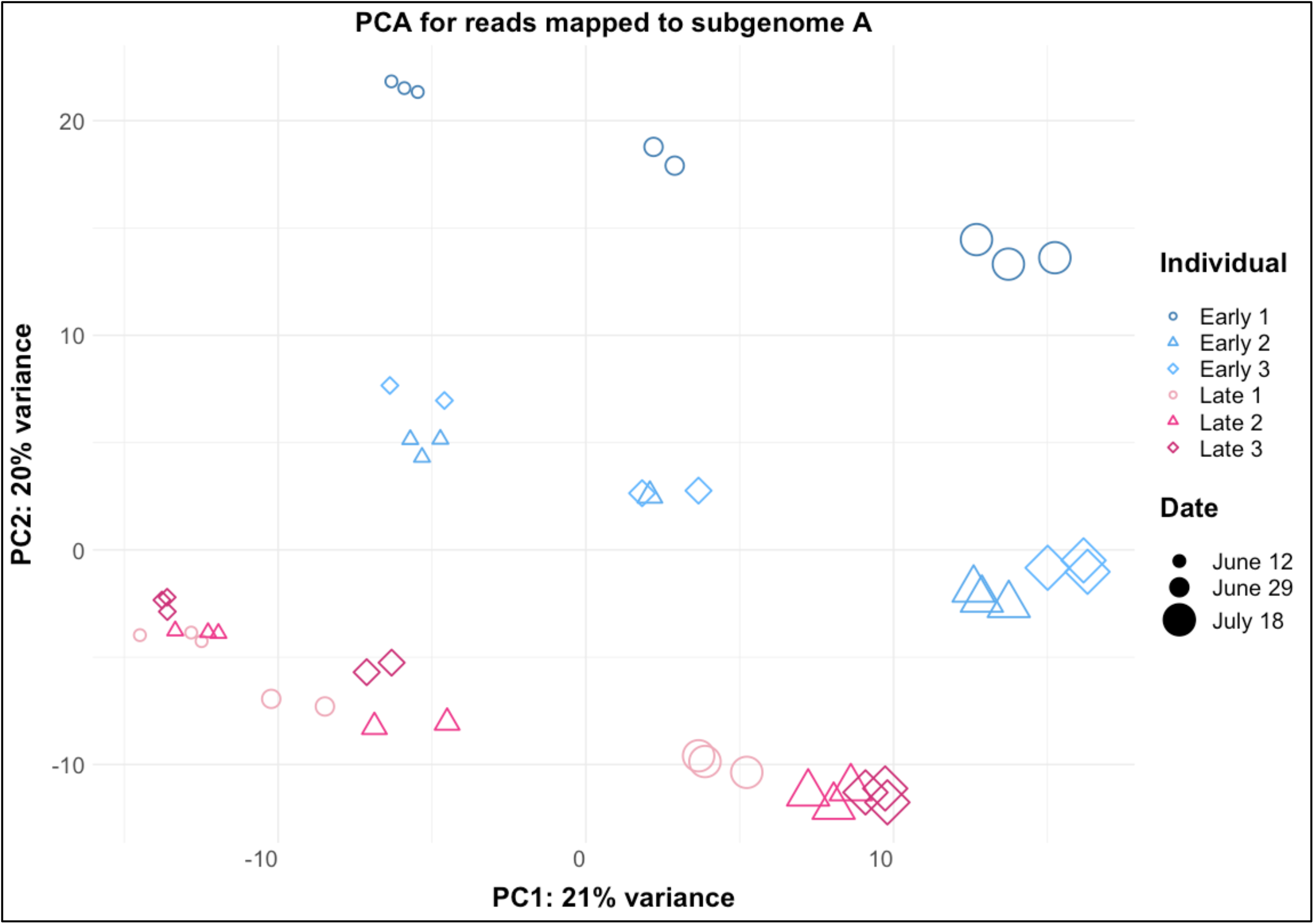
Principal Component Analysis (PCA) for three early-blooming and three late-blooming individuals suggests fundamental molecular differences between bloom groups. The PCA was produced with RNAseq data from 12 June 2019, 29 June 2019, and 18 July 2019 of EB and LB siblings’ apices in the floral position. PC1 appears to contain most of the variation seen from the different sampling dates while PC2 is at least partially driven by bloom group. RNAseq reads were mapped to subgenome A and two outliers were removed prior to producing the PCA.

On 12 Jun 2019, when all apices appeared vegetative (stage 1), 1371 genes were differentially expressed between the EB and LB bloom groups (**Figure 15A**). Approximately 13% (n = 183) of the DE genes for this date were located within the chromosome 4 QTL, despite this region only being approximately 3% of the genome (**Figure 15B**). The top 20 enriched gene ontology (GO) terms for genes more highly expressed in EBs (blue) and those more highly expressed in the LBs (pink) are shown in **Figure 16**. The top 20 terms support the notion that EBs were undergoing developmental processes mediated by jasmonic acid (JA), salicylic acid (SA), abscisic acid (ABA) and gibberellic acid (GA). The top 20 GO terms of genes more highly expressed in the LBs also included JA-related terms, as well as brassinosteroid (BR) and ethylene associated terms (**File S10**).

**Figure 15:**
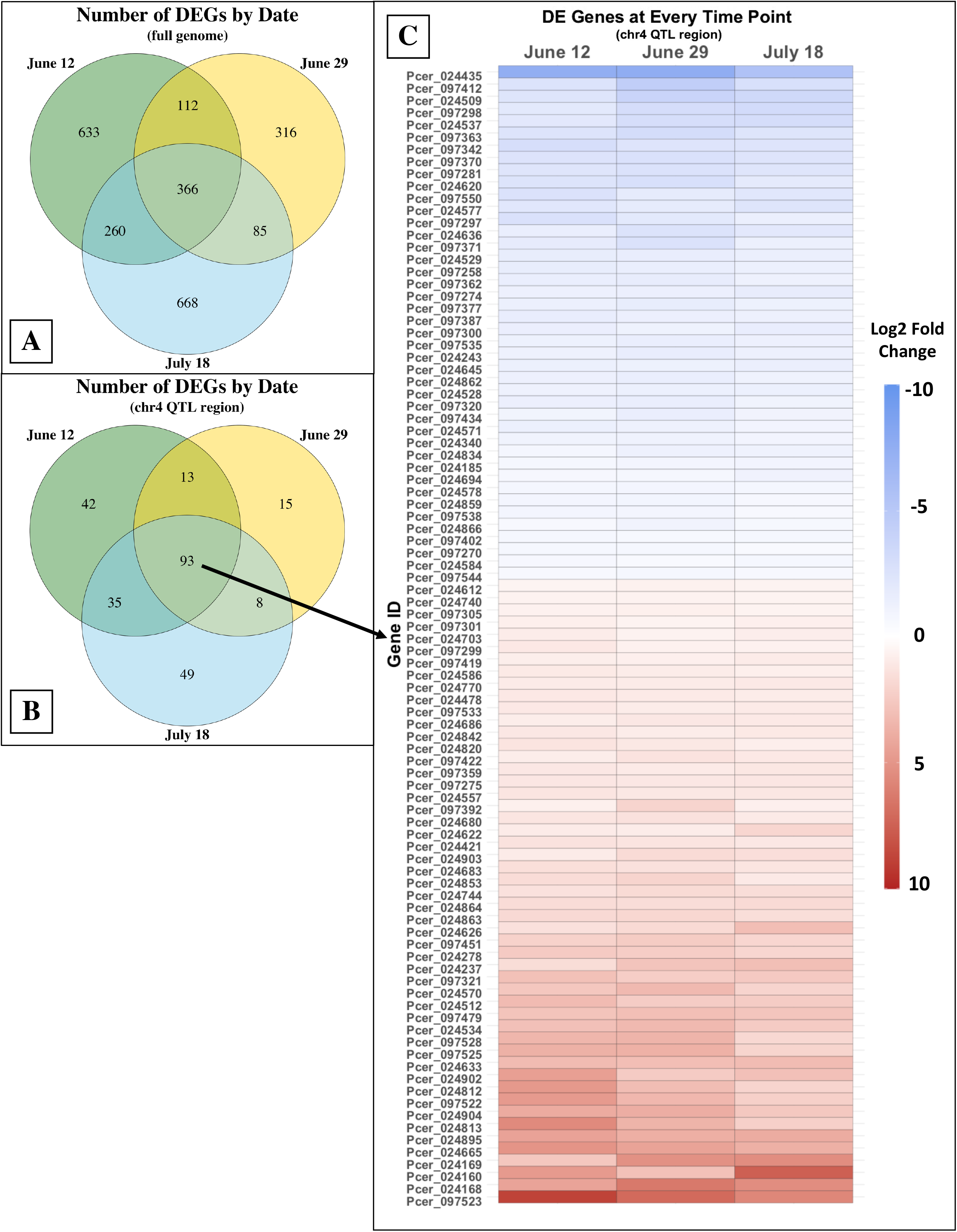
Venn diagrams and a heatmap illustrating the number of differentially expressed genes between early-blooming and late-blooming trees by date in the full genome and chromosome 4 QTL. A) Number of genes expressed by date throughout the genome, using subgenome A as a reference. B) Number of genes expressed by date only within the chromosome 4 QTL. C) A heatmap showing relative fold changes of DEGs at every date within the QTL. Blue shading represents genes more highly expressed in late-blooming individuals, while red shading represents genes more highly expressed in the early-blooming individuals. Gene numbers beginning in 097 were manually edited with Apollo (see Methods) and thus do not reflect relative order on the chromosome.

**Figure 16:**
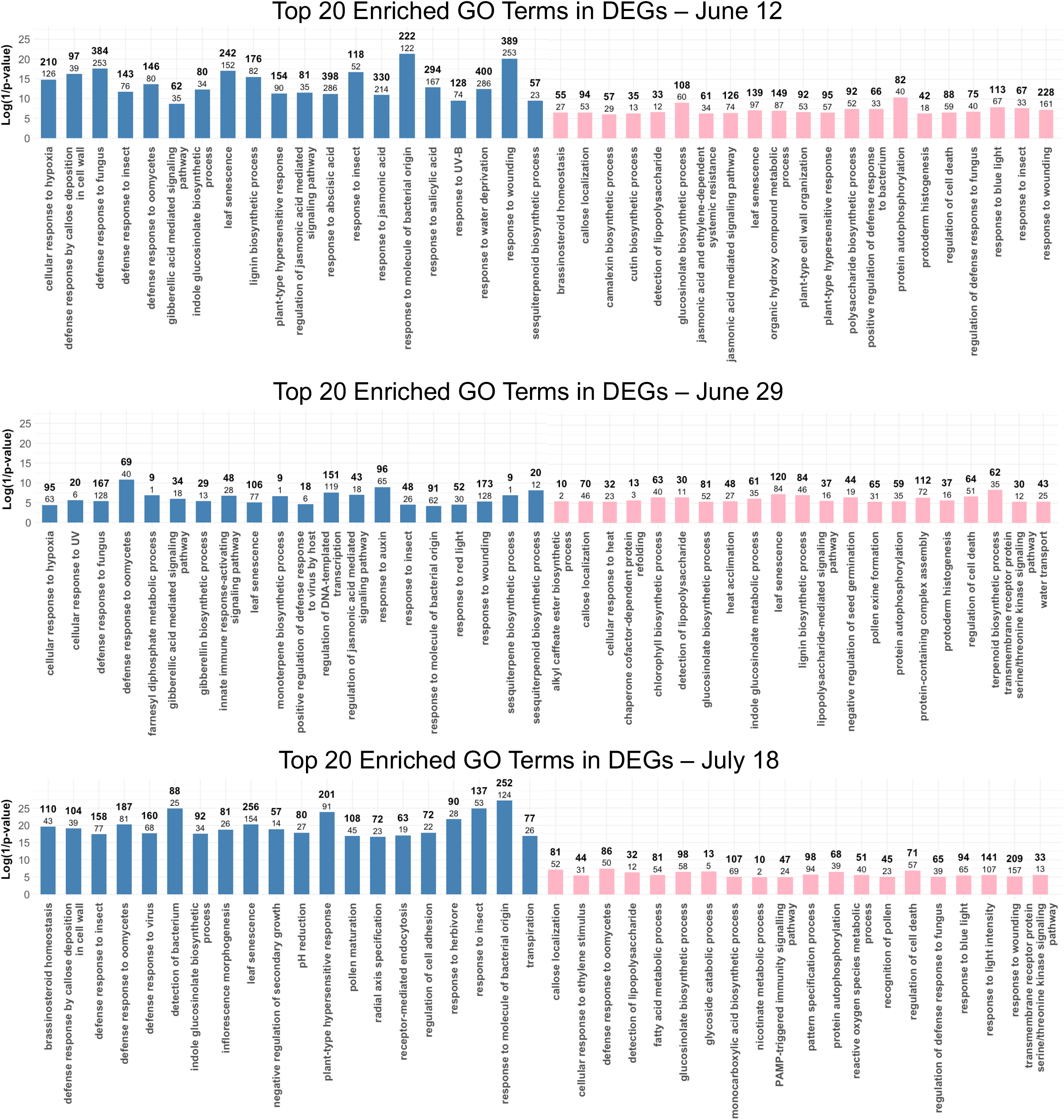
Results of the Gene Ontology enrichment analysis for each date’s comparison. The top 20 most enriched GO terms of genes more highly expressed in early-bloomers (blue) or late-bloomers (pink) are shown. The bolded number above each bar is the number of genes in the DEGs list with the respective GO annotation, while the number below is the amount expected based on the number annotated in the full database (DESeq2 object). The log value of the inverse p-value is plotted on the y-axis – thus, the taller the bar, the more significant the p-value according to a Fisher’s Exact Test. All p-values are below 0.05.

**Figure 17:**
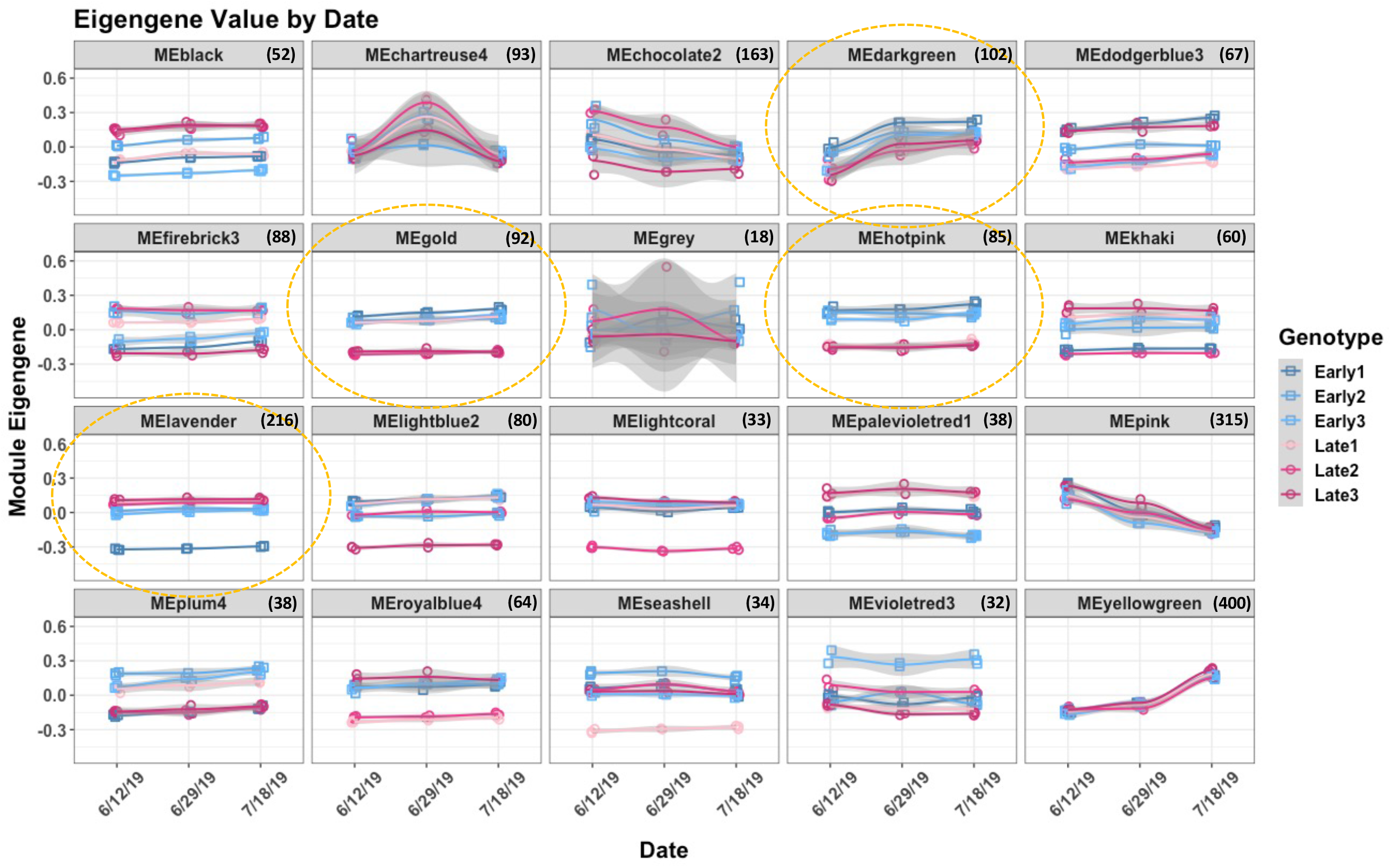
Module Eigengene (ME) expression trends for each module identified in the weighted gene correlation network analysis. MEs represent groups of genes with similar expression patterns and can help describe the differences in individuals. For example, the gene expression of the seashell module sets Late 1 apart from all individuals, while lightcoral sets Late 2 apart. The dotted gold circles draw attention to the hotpink, gold, darkgreen, and lavender modules, as all four separate the bloom groups to some degree. The number of genes housed in each module is given in parentheses next to the heading of each plot. A total of 2053 genes throughout the full genome were grouped into MEs. The hotpink module contains genes that most effectively, by far, separate the EB and LB siblings’ expression profiles.

On 29 Jun 2019, apices from both bloom groups were visibly still at stage 1. At this time, 879 genes were found to be differentially expressed between EBs and LBs, and nearly 15% of these DE genes (n = 129) were located within the bloom time QTL on chromosome 4 (**Figure 15A – B**). The top 20 GO terms for genes expressed at higher levels in the EBs implicate auxin-(IAA) and GA-mediated pathways at this stage of development (**Figure 16**) and GO terms among the top 50 implicate SA and ABA regulated processes. For genes more highly expressed in LBs, GO terms appear to be more related to basic metabolic and growth-related activities: for example, ‘lignin biosynthetic process,’ ‘regulation of cell death,’ and ‘protoderm histogenesis,’ might describe high rates of cell turnover.

On 18 Jul 2019, EB individuals showed a higher proportion of apices visibly making the transition from a vegetative to reproductive meristem (stage 2 and 3). 1379 DEGs between bloom groups were identified on this date, with 185 of these located within the chromosome 4 QTL (**Figure 15A – B**). The genes more highly expressed in the EBs support the morphological observations, with ‘inflorescence morphogenesis’ and ‘radial axis specification’ among the top 20 GO terms (**Figure 16**). For genes more highly expressed in the LBs, several of the top 20 terms are repeated from 29 June (e.g., ‘regulation of cell death,’ ‘protein autophosphorylation,’ and ‘callose localization’), perhaps suggesting fewer major developmental changes since 29 June. Markedly, 366 DEGs were common to all three dates, with 93 (25%) of these located in the chromosome 4 QTL (**Figure 15A – C**).

### Uniting genetic variation with differentially expressed genes

To obtain quality candidate genes for the basis of *k*, DEGs information was intersected with genetic variation. For smaller sources of variation (SNPs, INDELs < 30bp), we chose to focus on variation categorized as high effect (stop codon, frame shift, splice site changes), moderate effect (nonsynonymous amino acid changes, codon insertions/deletions), and certain modifier effects (upstream, 5’ UTR, and 3’ UTR) since these types would presumably affect protein function or gene regulation. For larger variants, only upstream and high effect structural variants were carefully considered. In total, 186 DEGs (for any date) in the QTL were predicted to have at least one of the above listed kinds of variation (**File S12**). 130 of these were expressed on 12 June 2019, 89 on 29 June 2019, 133 on 18 July 2019, and 62 were differentially expressed at all three dates.

### Weighted Gene Correlation Network Analysis

Next, we explored co-expression networks to identify groups of genes with expression profiles that distinguish the two bloom groups from one another. Post-filtering, 2053 genes were grouped into 19 modules ranging in size from 33 to 400 genes (excludes the grey module as these genes by definition do not fit well into any module; **Figure S13; File S11**). To visualize the expression profiles for each of these modules, the module eigengene values were plotted for each sample across the three time points (**Figure 17**). Several modules completely or partially separated the bloom time phenotypes, and the top 20 GO terms associated with these are provided in **Figure 18**. According to the GO enrichment analysis, the hotpink module (n = 85) suggested differences in terpene-based hormone synthesis, with genes in this module showing higher average expression in EBs compared to LBs on every date. The gold module (n = 92) showed similar biological terms but exhibited distinctions regarding abiotic stress (e.g., cellular response to water deprivation) and possibly hormone-regulated processes as it was enriched for strigolactone biosynthetic processes. It also displayed a similar expression pattern to that of hotpink with the exception of Late 1, whose expression profile was far less removed from the EBs compared to its fellow LBs. The darkgreen module (n = 102), which showed lower expression on 12 Jun compared to 29 Jun and 18 Jul for all samples, was enriched for terms related to auxin signaling and specialized tissue development. For example, suberin and lignin are likely components of the bud scales surrounding the apex, which continually form throughout the growing season. At any given date, EBs presented greater average expression of the genes in this module compared to LBs. Finally, LBs showed slightly higher expression of genes within the lavender module (n = 216), although Early 1 is somewhat of an outlier. This module was enriched for terms related to cell turnover, most notably ‘homeostasis of number of meristem cells.’ In agreement with the enriched GO terms, the hotpink module includes seven terpene synthases, a cytochrome P450 protein, and two AP2/ERF transcription factors. 15 genes in this module have some form of genetic variation, are differentially expressed, and are within the chr4 QTL. The gold module includes two MYB transcription factors, an ortholog of the transcription factor DEMETER, two cytochrome P450 proteins, and four terpene synthases. The darkgreen includes two MADS-box containing genes, three auxin-responsive proteins, and four cytochrome P450 proteins. For the lavender module genes, seven INSENSITIVE 1-ASSOCIATED RECEPTOR KINASES and a FANTASTIC FOUR-like protein were identified. The latter has been shown to regulate meristem size in arabidopsis (Wahl *et al*., 2010).

**Figure 18:**
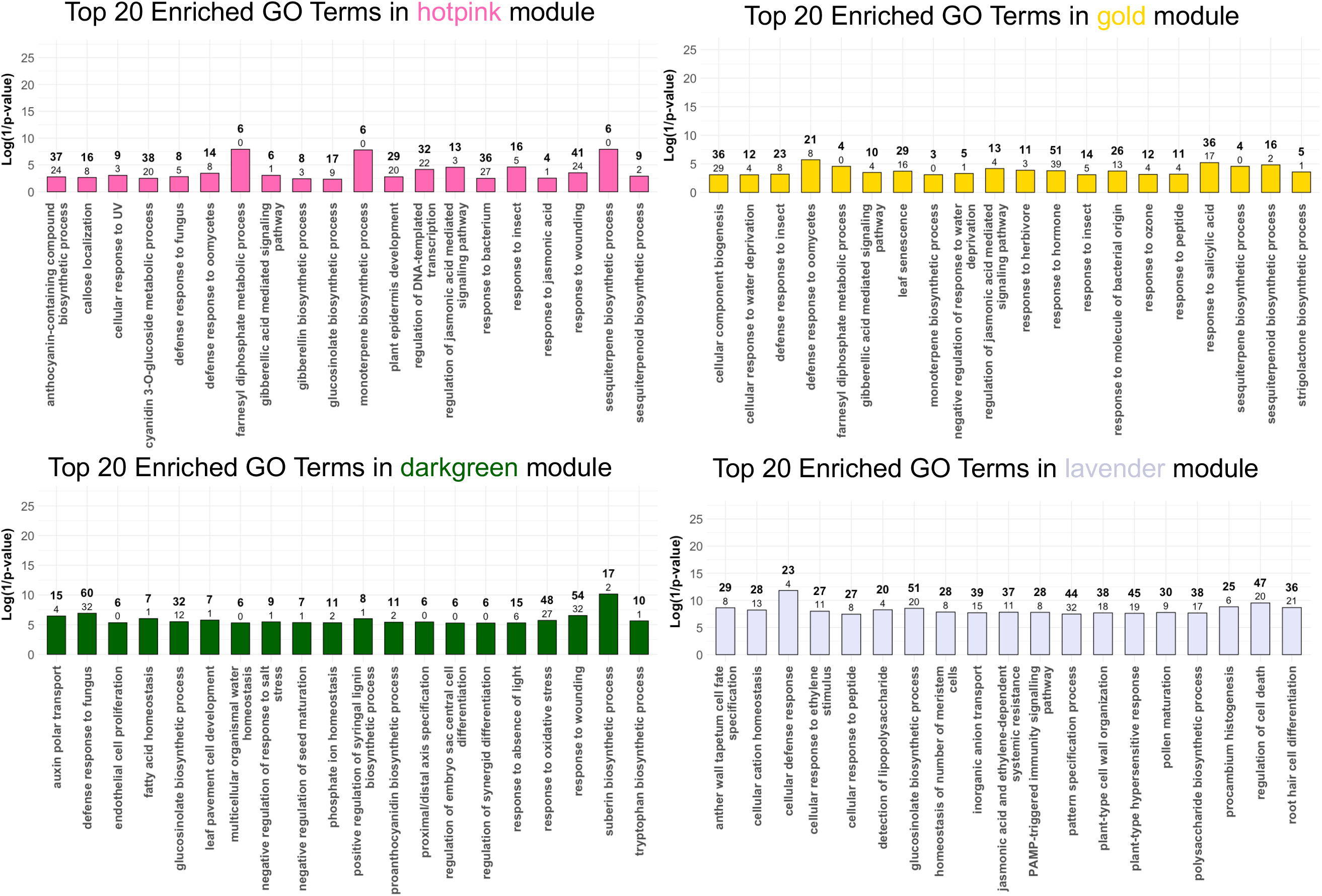
Results of the Gene Ontology enrichment analysis for select modules from the network analysis. The top 20 most enriched GO terms of genes found in the hotpink, darkgreen, gold, and lavender modules are shown. The bolded number above each bar is the number of genes in this module with the respective GO annotation, while the number below is the amount expected based on the number annotated in the full database (DESeq2 object). The log of the inverse p-value is plotted on the y-axis – thus, the taller the bar, the more significant the p-value according to a Fisher’s Exact Test. All p-values are below 0.05.

### Candidate genes for the basis of k

Gene candidates for the late bloom phenotype were prioritized based on location within the chr4 QTL, differential gene expression, amount and type of genetic variation, co-expression with genes in modules that distinguished early- and late-blooming individuals (**Figure 17**), and their predicted gene function (**Table 1**). According to the PCA, bloom groups were separable transcriptionally by 12 June despite all apices being vegetative, suggesting the *k* haplotype could impact phenology even before floral initiation. Thus we reasoned the greatest chance of capturing the activity of gene(s) behind the basis of *k* would be on the earliest date, 12 June. Genes that were differentially expressed on at least 12 June (i.e., expressed only on 12 June, 12 June + [29 June or 18 July] or at all time points) were thus favored. Eight of the 13 highest-ranked candidates were co-expressed with genes in the hotpink module as it best delineated the bloom groups.

The fine-mapping results for a chromosome 4 bloom time QTL discovered in a population developed from sweet cherry (*P. avium*) cultivars ‘Regina’ and ‘Garnet’ were also taken into account when considering candidates (Branchereau *et al*., 2022). This QTL was associated with a delay in bloom time of nearly 4 days (Dirlewanger *et al*., 2012). The peak position of the chr4 sour cherry QTL was barely outside the 95% confidence interval of the one identified the sweet cherry, but since their confidence intervals overlapped, they were previously considered the same QTL (Cai *et al*., 2018). We therefore identified the syntenic region of the fine-mapped sweet cherry QTL on sour cherry chromosome 4A (approx. Pcer_chr4A: 9717998 - 9802438) and surveyed genes within this region for differential expression for any dates sampled in the current study. However, there were none – suggesting these QTL are distinct from one another, and the effect of *k* may be due to separate gene(s).

Among the topmost ranked candidates are several terpene synthases and B3 domain-containing transcription factors. The terpene synthases are of considerable interest since hormones known to be involved in several phenological and developmental transitions are terpene-based (e.g., GA, ABA) (Bradley and Crane, 1960; Taiz and Zeiger, 2010; Vimont *et al*., 2019; Villar *et al*., 2020). One terpene synthase (Pcer_097523-RA) has a single base pair deletion anticipated to induce a frame-shift in the *k* allele – consistent with this prediction, the expression has been reduced to nearly nothing in the late-blooming individuals. Several of the B3 domain-containing transcription factors are orthologs of arabidopsis AT5G66980, which has a strikingly floral-specific expression pattern and belongs to a B3 domain-containing subfamily of proteins called REPRODUCTIVE MERISTEM (REMs) (Mantegazza *et al*., 2014). Like other members of this subfamily, these three candidates exist in a tandem array of six genes within an 18 kb region (Pcer_chr4A: 13481626 - 13500091). However, the predicted protein sequence of the fourth reads through one stop codon and was initially missed during annotation (Pcer_024686-RA).

## DISCUSSION

Various studies have tried to identify connections between early floral development and final bloom time in fruit trees. In 1984 and 1985, Raseira and Moore grouped nine peach cultivars into low (< 500 chilling hours below 7.2 °C), medium (550 – 750), and high (800 – 1050) chill requirements and assessed floral bud initiation times. Between the two years, it was extremely variable: in 1984, the low-chill group was the first to initiate floral buds, whereas the high-chill group initiated theirs first in 1985 (Raseira and Moore, 1987). In 1989, Hoover *et al*. tracked floral development in four *Malus* × *domestica* cultivars and found cultivar-specific rates of organogenesis, but no clear association between initiation and bloom time (McArtney *et al*., 2001). Fadón *et al*. (2018*b*) investigated early floral development in five sweet cherry cultivars and noted asynchronous progression among the genotypes for all three years, but that a common stage was reached by dormancy – albeit at different times. More recently, Goeckeritz *et al*. described the pre-dormancy development of six apple accessions – four species total – and showed a wide range of stages at approximate dormancy (Goeckeritz *et al*., 2023*a*).

Aside from other limitations of each study – including the scale at which flowers were examined (e.g., cellular level vs macrodissections) – the wide genetic variation among sampled trees was not addressed (Goeckeritz *et al*., 2023*a*; Raseira and Moore 1987, Hoover *et al*.1989, Fadon *et al*., 2018*b*). Such genetic variation likely clouds our ability to identify subtle changes in phenology between genotypes, which could explain why an association for floral initiation and final bloom time has yet to be found. Many genes affect bloom time, so it is quite probable that several mechanisms are confounded when contrasting diverse materials. However, confronting this issue is easier said than done as research orchards are expensive to maintain and access to segregating populations is rare. Additionally, ‘domesticated’ fruit trees are often not far genetically removed from related wild populations and pedigrees are unknown. The work detailed here takes advantage of the genetic relationships in a population of sour cherry created at the Michigan State University Clarksville Research Center. We built on previous work identifying a major QTL on linkage group 4, a hotspot for bloom time regulation in *Prunus* (Cai *et al*., 2018). While controlling for genetic relatedness, we sought to identify the changes in flower development associated with one particular haplotype, *k*, which additively delays bloom time by three days. Guided by these phenological assessments, we present strong candidate genes for future functional work.

Other studies that searched for candidate genes within *Prunus* chromosome 4 QTL focused on dormancy, which is indisputably critical to bloom time in Rosaceous fruit trees (Romeu *et al*., 2014; Dirlewanger 2015; Castede *et al*., 2015; Branchereau *et al*., 2022). However, doing so fails to address the developmental events that have occurred up until that point. Furthermore, long generational times and few genomic resources for *Prunus* led to these studies being limited in their scope and narrowly-focused on homologs known to affect flowering time in annual species. For instance, a *SQUAMOSA PROMOTER-BINDING LIKE* gene (*SPL5*), *LFY*, *SUPPRESSOR OF OVEREXPRESSION OF CONSTANS* (*SOC1*), and *PHOTOPERIOD INSENSITIVE EARLY-FLOWERING* (*PIE1*) have all been cherry-picked as candidates over the years (Moon *et al*., 2003; Noh and Amasino, 2003; Silva *et al*., 2005; Wu and Poethig, 2006; Dirlewanger *et al*., 2012; Castede *et al*., 2015). Fortunately, resources are rapidly being generated as genome sequencing becomes routine (Verde *et al*., 2017; Jiang *et al*., 2019; Wang *et al*., 2020; Yi *et al*., 2020; Cao *et al*., 2022; Goeckeritz *et al*., 2023*b*). Recently, researchers successfully fine-mapped a chromosome 4 QTL in *P. avium* (sweet cherry) to just 12 genes within a 68 kb region in the ‘Regina’ genome (Branchereau *et al*., 2022). Alas, this locus does not overlap with any of the candidate genes proposed in previous studies – a cautionary tale for choosing candidate genes based solely on homology (Baxter, 2020).

Our morphological and quantitative data show EBs surpass their LB siblings as soon as floral development begins, and that their lead is maintained even up until bloom the following spring. Assessing molecular distinctions as close to the time of divergence as possible increases the chances of capturing the expression of gene(s) driving the phenotypic differences while controlling for confounding effects of changes that may have already occurred. RNA-sequencing data indicated that, despite their morphological similarities, gene expression profiles were fundamentally different between the bloom groups at all dates sampled – suggesting the effect of *k* may act even before the apex has committed to the floral transition. Although the floral development staging system designed in the present work was useful in characterizing histological variation between EBs and LBs, the molecular differences identified via RNA-sequencing underscore the limitations of this method. These limitations may be the reason that pre-dormancy morphological variation appears to expand during organogenesis and shrink as the flowers enter dormancy in the two years this developmental window was observed.

Based on the gene expression data and subsequent GO term analyses, the EB and LB individuals fine tune their hormone balances during the vegetative to reproductive phase transition, which is in accordance with observations in sweet cherry and apple (Li *et al*., 2018; Villar *et al*., 2020). Examination of the GO terms found in upregulated DEGs as EBs proceeded from 12 June to 29 June included ‘response to ABA’ and ‘response to IAA’ paired with ‘vegetative to reproductive phase transition,’ lending support to the positive influences ABA and IAA may have during this transition in fruit trees (Villar *et al*., 2020). The LBs show a strikingly diverse profile – in addition to ABA and IAA, they show enrichment terms related to BR, JA, SA, CK, and GA, the latter of which is robustly connected to maintenance of the vegetative phase in fruit trees (Bradley and Crane, 1960; Zhang *et al*., 2019; Gottschalk *et al*., 2021). When including GO information for co-expression modules, a consistent theme separates the bloom groups: terpene synthesis. Terpene compounds are greatly diversified in plants and take part in everything from biotic defense responses to phytohormone synthesis (Tholl, 2006). Gibberellins constitute a group of tetracyclic diterpenes (C_20_), ABA is synthesized from a carotenoid precursor and classified as a tetraterpenoid (C_40_), and BR is made from the triterpene (C_30_) squalene (Taiz and Zeiger, 2010). Terpene synthases, the enzymes responsible for these chemical reactions, may form species-specific paralagous gene clusters with tissue-specialized gene expression (Tholl, 2006; Mantegazza *et al*., 2014; Jia *et al*., 2022).

The hotpink module identified in the network analysis (n = 85) most effectively separated the bloom groups and contained seven terpene synthases as well as several hormone-related GO terms (**Figure 18**). Variable GA responses and/or synthesis as the basis of *k* is an interesting possibility; GA is linked to the floral transition, and its relationship with ABA is a key indicator of dormancy depth (Goeckeritz and Hollender, 2021). In the two seasons of forcing experiments, it was shown that EBs responded to warm temperatures first; this implies a shallower state of dormancy and the possibility that *k* may affect temperature responses as well. In 2019 – 2020, we attempted to saturate the chilling response for both bloom groups by placing branches in a 5 °C chamber before forcing (CP = 91.5). However, as shown in **Figure 11**, this only served to further separate these individuals as additional chill caused faster growth rates for the EBs compared to LBs. This trend is most striking at later stages of budbreak (BBCH 54), which is also observed in the gradual divergence of the pistil’s growth (**Figure 10**). The apparent differences in temperature-sensing mechanisms among the bloom groups fits well with GA’s known involvement in these pathways (Stavang *et al*., 2009). The apparent paradox that higher GA levels negatively regulate the floral transition in rosaceous fruit trees but positively regulate dormancy transitions and spring growth could be explained by their spatiotemporal context. For example, in arabidopsis, GA signaling first encourages the floral transition but then later prevents flower formation in the lateral primordia of the inflorescence (Yamaguchi *et al*., 2014). Taken together, these results advise a closer examination of terpene-based hormone synthesis as a potential means of regulating bloom time in sour cherry.

Also among the promising candidate genes in **Table 1** are four differentially expressed B3 domain-containing transcription factors, two of which are co-expressed with genes in the hotpink module. *AUXIN RESPONSE FACTORS* (*ARFs*), *ABSCISIC ACID INSENSITIVE 3* (*ABI3*), *HIGH LEVEL EXPRESSION OF SUGAR INDUCIBLE* (*HSI*), *RELATED TO ABI3/VP1* (*RAV*), and *REPRODUCTIVE MERISTEM* (*REM*) genes make up five subfamilies within the B3 domain-containing superfamily (Romanel *et al*., 2009). The candidates noted here all share closest homology with the *REM* subfamily, which coincidentally has the most sequence divergence among this family and the highest rate of expansion (Franco-Zorrilla *et al*., 2002; Romanel *et al*., 2009). Usually, they exist in a tandem array, with *REM* gene clusters showing more sequence similarity within themselves than between clusters (Romanel *et al*., 2009). As their name implies, a survey of the expression patterns in arabidopsis shows they are tissue-specific – at times, only expressed in a few cells – with various roles in flower development (Mantegazza *et al*., 2014; Caselli *et al*., 2019). However, due to their duplicative nature, functional redundancy and tight genetic linkage has made identification of specific phenotypes difficult (Mantegazza *et al*., 2014). The best functionally-characterized member of the *REM* subfamily is *VERNALIZATION 1* (*VRN1*), a gene required for the silencing of *FLC* and the accelerated flowering time seen after exposing arabidopsis to low temperatures for several weeks (Chandler *et al*., 1996; Noh and Amasino, 2003). More recently, REM17 was shown to be a positive effector of flowering time that is negatively targeted by FLC and SVP1 and positively targeted by SOC1 after vernalization (Richter *et al*., 2019). Furthermore, REM16 was identified as a flowering time regulator through its interaction with ACTIN DEPOLYMERIZING FACTOR 1 (ADF1); REM16 was shown to directly bind to both the *SOC1* and *FT* promoters to enhance their expression (Yu *et al*., 2020). The B3 domain-containing/REM proteins associated with the late-blooming phenotype in this sour cherry population are therefore good candidates for further functional analysis.

As discussed earlier, there is evidence that genes regulating bloom time in one *Prunus* species may regulate bloom time in another (Jimenez *et al*., 2010; Calle *et al*., 2021). Therefore, we compared the expression of candidate genes for an overlapping bloom time QTL in sweet cherry to the sour cherry chr4 QTL DEGs identified in the present work. Previously, both QTL were considered to be the same (*qP-BD4.1^m^*); here, the syntelogs of the genes in sweet cherry showed no clear expression patterns between our sour cherry bloom groups. It is possible we were unable to capture the proper temporal period for the differential expression of these genes. However, it should be noted that the number of DEGs between the EBs and LBs was especially enriched in the chr4 QTL. Therefore, it is reasonable to suspect more than one major regulator of bloom time is present on *Prunus* chromosome 4. Further supporting multiple regulators is the finding that *k* is derived from subgenome A, a *P. fruticosa*-like subgenome (Bird *et al*., 2022; Goeckeritz *et al*., 2023*b*). Given the contrasting selective pressures in the native habitats of *P. fruticosa* and *P. avium*, we now have incentive to believe the bloom time variation in these species is conferred by separate genes. In addition to *k*, other alleles in sour cherry have unique potential to additively delay bloom in other *Prunus* species since some accessions of sour cherry are the latest bloomers of the entire genus (Iezzoni personal communication; Cai *et al*., 2018).

In conclusion, our assessment of flower development in early- and late-blooming sour cherry siblings from pre-initiation to anthesis suggest comparisons starting from dormancy may be limiting our understanding of flowering time in fruit trees. The sectioning data clearly show developmental differences as soon as floral initiation, and transcriptional evidence supports molecular distinctions well before this. The pistil measurements support pre-dormancy discrepancies as well, and the forcing experiments suggest contrasts in temperature perception. It is unclear whether the stage of development at dormancy entrance may affect transitions as this question remains largely unexplored. Finally, both the forcing experiments and pistil data support variable rates of growth between the bloom groups as flowers resume development in spring. We gathered this data to identify the specific effect(s) the gene(s) within a major chr4 QTL have on final bloom time. In the process, we identified promising candidate genes underlying the late-blooming phenotype of haplotype *k* for future functional study. As the climate continues to change, phenological data like these are needed to anticipate how long-lived crops such as fruit trees will respond to environmental perturbations. In short, our work suggests there is indeed more to flowering than those *DAM* genes.

## SUPPLEMENTARY INFORMATION

**Table S1**: The identities of each tree in the research orchard used for the present study, their *k* dosage, relative bloom times, identification number for their bloom group, and bloom dates for several years

**Table S2**: Dates flowers were collected for pistil measurements

**Table S3**: Dates flowers were collected for histological assessments

**Table S4**: Dates branches with floral buds were collected for the forcing experiments

**Table S5**: Dates flowers were collected for the H_2_O_2_ measurements

**Figure S1:** Anther sections of one early-blooming and one late-blooming sibling on the same day in late February 2018 suggest developmental differences occur prior to dormancy exit

**Figure S2:** Sections of early-blooming and late-blooming siblings on 14 June 2018

**Figure S3:** Sections of early-blooming and late-blooming siblings on 8 August 2018

**Figure S4:** Sections of early-blooming and late-blooming siblings on 20 September 2018

**Figure S5:** Sections of early-blooming and late-blooming siblings on 30 October 2018

**Figure S6:** Sections of early-blooming and late-blooming siblings on 27 November 2018

**Figure S7:** Sections of early-blooming and late-blooming siblings on 27 December 2018

**Figure S8:** Sections of early-blooming and late-blooming siblings on 21 February 2019

**Figure S9:** Sections of early-blooming and late-blooming siblings on 18 March 2019

**Figure S10:** Sections of early-blooming and late-blooming siblings on 5 April 2019

**Figure S11:** Sections of early-blooming and late-blooming siblings on 15 April 2019

**Figure S12:** Association between H_2_O_2_ content, a reactive oxygen species (ROS), and the temperature at the time of collection

**Figure S13:** The hierarchal clustering of gene expression and module groupings indicate strong clusters were identified during the WGCNA analysis

**File S1**: Raw pistil measurements for 2018-2019

**File S2**: Raw pistil measurements for 2019-2020

**File S3**: Raw pistil measurements for 2021-2022

**File S4**: Data for forcing experiments in 2019-2020

**File S5**: Data for forcing experiments in 2021-2022

**File S6**: Data for H_2_O_2_ quantification in flower buds in 2019-2020

**File S7**: Haplotype and subgenome information for deducing *k* is a unique allele derived from subgenome A

**File S8**: Lists of filtered genetic variation predicted in the SNP and SV analyses that are unique to individuals with 2*k*

**File S9**: Lists of differentially expressed genes in early- and late-blooming siblings’ floral-positioned apices, including their fold changes

**File S10**: Top 50 GO terms for genes upregulated and downregulated in select comparisons **File S11**: Genes found in each WGCNA module and the top 50 GO terms enriched in each network

**File S12**: Candidate genes for the basis of the *k* haplotype, including gene names, putative functions, predicted genetic variation, WGCNA modules, and more

## ACKNOWLEDGEMENTS

The authors would like to thank the staff at the Michigan State University Clarksville Research Center – especially Dan, Anderson, Jerry, and Tina – for maintenance and preservation of materials. The authors are grateful to the College of Agriculture and Natural Resources’ Statistical Consulting Center, Dr. Addie Thompson, Dr. Miranda Haus, and Andrea Kohler for guidance with statistical analyses. Dr. Joseph Hill assisted in troubleshooting ROS extraction protocols. Dr. Robert VanBuren and Dr. Ben Mansfeld kindly provided advice for differential gene expression and network analyses, and Kathleen Rhoades provided sequencing data for cultivars ‘Balaton’ and ‘Surefire.’ Asyraf Azmi was instrumental in assisting with field collections and experiments, and Dr. Alicia Withrow was essential to all microscopy work.

## AUTHOR CONTRIBUTIONS

CZG, CAH, RG, AI: conceptualization; CZG: methodology; CZG: formal analysis; CZG, CG: investigation; CZG: writing – original draft; CZG, CG, CAH, RG, and AI: review & editing; CZG: visualization; CAH and AI: funding acquisition.

## DATA AVAILABILITY

The datasets, code, and scripts supporting the conclusions of this article are available at https://github.com/goeckeritz/sourcherry_bloomtime. Raw RNA and DNA sequencing data can be found in NCBI’s Sequence Retrieval Archive (SRA) under BioProject number PRJNA1036689.

